# Maximum Entropy Framework For Inference Of Cell Population Heterogeneity In Signaling Networks

**DOI:** 10.1101/137513

**Authors:** Purushottam D. Dixit, Eugenia Lyashenko, Mario Niepel, Dennis Vitkup

**Affiliations:** Department of Systems Biology, Columbia University; Department of Systems Biology, Harvard Medical School; Department of Biomedical Informatics, Columbia University; Center for Computational Biology and Bioinformatics, Columbia University

## Abstract

Predictive models of signaling networks are essential tools for understanding cell population heterogeneity and designing rational interventions in disease. However, using network models to predict signaling dynamics heterogeneity is often challenging due to the extensive variability of signaling parameters across cell populations. Here, we describe a **M**aximum **E**ntropy-based f**R**amework for Inference of heterogeneity in **D**ynamics of s**I**g**A**ling **N**etworks (MERIDIAN). MERIDIAN allows us to estimate the joint probability distribution over signaling parameters that is consistent with experimentally observed cell-to-cell variability in abundances of network species. We apply the developed approach to investigate the heterogeneity in the signaling network activated by the epidermal growth factor (EGF) and leading to phosphorylation of protein kinase B (Akt). Using the inferred parameter distribution, we also predict heterogeneity of phosphorylated Akt levels and the distribution of EGF receptor abundance hours after EGF stimulation. We discuss how MERIDIAN can be generalized and applied to problems beyond modeling of heterogeneous signaling dynamics.

## Introduction

Signaling cascades in genetically identical cells often respond in a heterogeneous manner to extracellular stimuli (Raj and van Oudenaarden 2008). This heterogeneity arises largely due to cell-to-cell variability in biochemical signaling parameters such as reaction propensities and chemical species abundances (Albeck, Burke et al. 2008, Spencer, Gaudet et al. 2009, Meyer, D’Alessandro et al. 2012, Llamosi, Gonzalez-Vargas et al. 2016, Kallenberger, Unger et al. 2017). This variability in parameters can have important functional consequences, for example, in fractional killing of cancer cells treated with chemotherapeutic drugs (Albeck, Burke et al. 2008, Spencer, Gaudet et al. 2009). Therefore, the knowledge of the distribution over parameters is essential to understanding phenotypic heterogeneity in cell populations.

Several experimental techniques such as flow cytometry (Wu and Singh 2012), immunofluorescence (Wu and Singh 2012), and live cell assays (Meyer, D’Alessandro et al. 2012) have been developed to investigate the variability of biochemical species abundances. However, it is often difficult to estimate the distribution over biochemical parameters from these experimental measurements. The reasons for this challenge are primarily threefold. First, parameters such as protein abundances and reaction propensities vary substantially across cells in a population (Raj and van Oudenaarden 2008). For example, previous studies have reported the coefficients of variation of protein abundances in the range 0.1 – 0.6 (Niepel, Spencer et al. 2009). This would make the effective rates of signaling reactions vary substantially between cells as well (Chung, Sciaky et al. 1997, Meyer, D’Alessandro et al. 2012). Second, the multivariate parameter distribution can potentially have a complex shape. For example, as is often the case in single cell data, gene expression, and thus cellular abundance, of key signaling proteins may exhibit multimodality (Shalek, Satija et al. 2013). Finally, single cell measurements are typically not sufficient to uniquely infer the underlying parameter variability; the problem usually referred as parameter non-identifiability (Banks 2012).

Over the last decade, several computational methods have been developed to estimate the joint distribution over parameters consistent with experimentally measured cell-to-cell variability in biochemical species (Waldherr, Hasenauer et al. 2009, Hasenauer, Waldherr et al. 2011, Zechner, Ruess et al. 2012, Hasenauer, Hasenauer et al. 2014, Zechner, Unger et al. 2014, Loos, Moeller et al. 2018, Waldherr 2018). Most of these methods circumvent the ill-posed inverse problem of estimation of the parameter distribution (Banks 2012) by making specific *ad hoc* choices about the shape of the distribution. For example, Hasenauer et al. (Hasenauer, Waldherr et al. 2011, Hasenauer, Hasenauer et al. 2014) (see also (Waldherr, Hasenauer et al. 2009, Loos, Moeller et al. 2018)) approximate the parameter distribution as a linear combination of predefined distribution functions. Waldherr et al. (Waldherr, Hasenauer et al. 2009) approximate the parameter distribution using Latin hypercube sampling. Similarly Zechner et al. (Zechner, Ruess et al. 2012, Zechner, Unger et al. 2014) assume that the parameters are distributed according to a log-normal or a gamma distribution. Consequently, it may be difficult for these previously developed approaches to infer a complex multivariate parameter distribution of an unknown shape.

Building on our previous work (Dixit 2013, Eydgahi, Chen et al. 2013), we developed **MERIDIAN:** a **M**aximum **E**ntropy-based f**R**amework for Inference of heterogeneity in **D**ynamics of s**I**gn**A**ling **N**etworks. Instead of enforcing a specific functional form of the parameter distribution *a priori*, MERIDIAN uses data-derived constraints to derive it *de novo*. The maximum entropy principle was first introduced more than a century ago in statistical physics (Dixit, Wagoner et al. 2018). Among all candidate distributions that agree with imposed constraints, the maximum entropy principle selects the one with the least amount of over-fitting. Maximum entropy-based approaches have been successfully applied previously to a variety of biological problems, including protein structure prediction (Weigt, White et al. 2009), protein sequence evolution (Mora, Walczak et al. 2010), neuron firing dynamics (Schneidman, Berry et al. 2006), molecular simulations (Dixit, Jain et al. 2015, Tiwary and Berne 2016), and dynamics of biochemical reaction networks (Dixit 2018).

Following a description of the key ideas behind MERIDIAN, we illustrate its performance using synthetic data on a simplified model of growth factor signaling. Next, we use the framework to study the heterogeneity in the signaling network leading to phosphorylation of protein kinase B (Akt). Epidermal growth factor (EGF)-induced Akt phosphorylation governs key intracellular processes (Manning and Toker 2017) including metabolism, apoptosis, and cell cycle entry. Due to its central role in mammalian signaling, aberrations in the Akt pathway are implicated in multiple diseases (Herbst 2004, Manning and Toker 2017). We apply MERIDIAN to infer the distribution over signaling parameters using experimentally measured levels of phosphorylated Akt (pAkt) and cell surface EGFR (sEGFR) in MCF10A cells (Soule, Maloney et al. 1990) following EGF stimulation. We then demonstrate that the obtained parameter distribution allows us to accurately predict the heterogeneity in single cell pAkt levels at late time points, as well as the heterogeneity in cell surface EGFRs in response to EGF stimulation. Finally, we discuss the generalizations of the framework to study problems beyond modeling heterogeneity in signaling networks.

## Results

### Outline of MERIDIAN

We consider a signaling network comprising *N* chemical species whose intracellular abundances we denote by 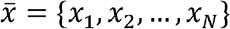. We assume that the molecular interactions among the species are described by a system of ordinary differential equations

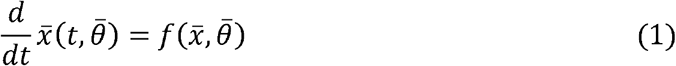

where 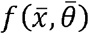 is a function of species abundances 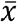. Here, 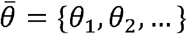 is a vector of parameters that describe the dynamics of the signaling networks. We denote by 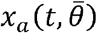 the solution of equations (1) for species “*a*” at time *t* with parameters 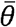.

Our computational approach is illustrated in Figure 1. We use experimentally measured cell-to-cell variability of protein species “*a*” at multiple experimental conditions, for example, several time points, (illustrated by histograms in Figure 1) to constrain the parameter distribution 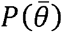. Specifically, we first quantify the experimentally measured cell-to-cell variability by estimating bin fractions *ϕ_ik_*. In our notation, the index *i* specifies the experimental condition (measurement time, measured species, input conditions, etc.) and *k* indicates bin number. Every distinct dynamical trajectory 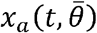 (illustrated by red and blue curves in Figure 1) generated by specific parameter values 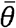 passes through a unique set of bins (red curve through red bins and blue curve through blue bins in Figure 1) at multiple experimental conditions. Using MERIDIAN, we find a corresponding probability distribution over signaling parameters 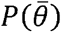 such that the distribution over trajectories 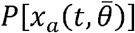 is consistent with all experimentally measured bin fractions. Below, we present our development to derive the functional form of 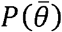 that is consistent with these constraints.

**Figure 1.**
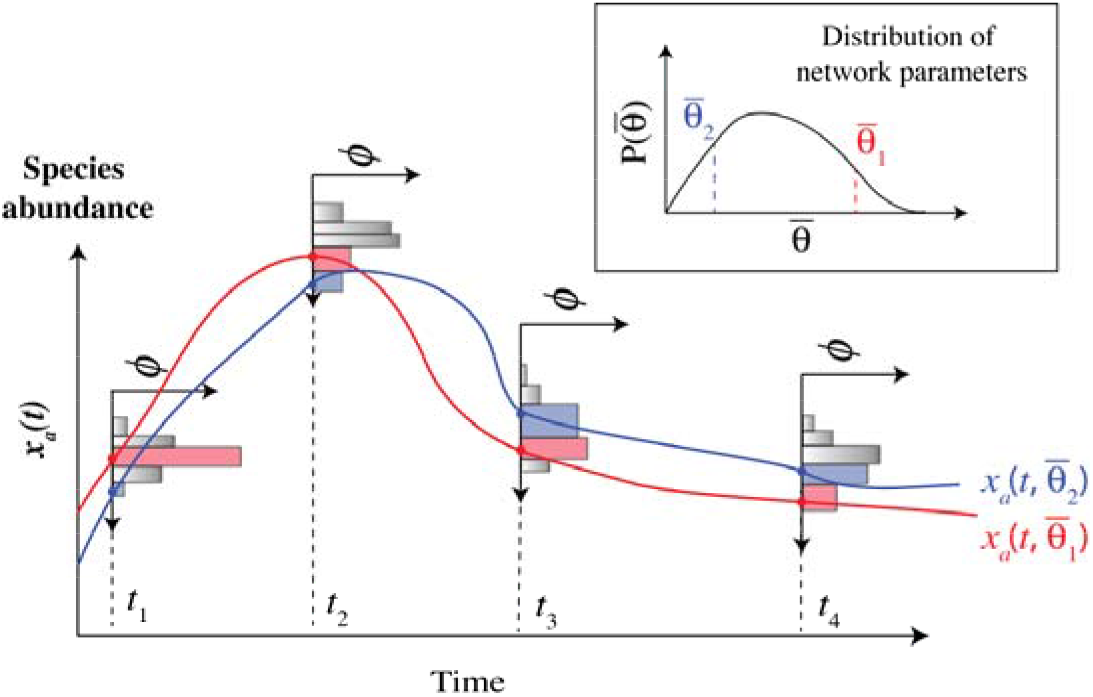
Illustration of the MERIDIAN inference approach. Cell-to-cell variability in protein “*a*” is measured at four time points *t*_1_, *t*_2_, *t*_3_, and *t*_4_. From the single cell data, we determine the fraction *ϕ_ik_* of cells that populate the *k*^th^ abundance bin in the *i*^th^ experiment. The histograms show *ϕ_ik_* at multiple experimental conditions. We find 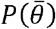 with the maximum entropy while requiring that the corresponding distribution 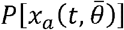 over trajectories of 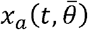 simultaneously reproduces all bin fractions.

### Derivation of 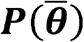 using MERIDIAN

For simplicity, we consider the case when the distribution of cell-to-cell variability in one species *x_a_* is available only at one time point *t* (for example, *t* = *t*_1_ in Figure 1). We denote by 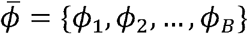 the fraction of cells whose experimental measurement of *x_a_* lies in individual bins. Here, given that we are considering only one experimental condition, for brevity we use only one index to indicate the bin fractions. Below, we first assume that there are no experimental errors in determining 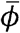. Later, we demonstrate how to incorporate known experimental errors both in the inference procedure and in making predictions using 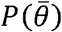.

Given a parameter distribution 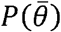 the predicted fractions 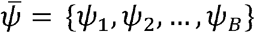 can be obtained as follows. Using Markov chain Monte Carlo (MCMC), we generate multiple parameter sets from 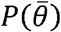. For each sampled set of parameters 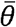, we solve equations (1) and find 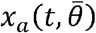, i.e. the predicted value of the abundance at time *t*. Then, using the samples from the ensemble of trajectories, we estimate *ψ_k_* as the fraction of sampled trajectories where 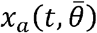 passed through the *k*^th^ bin. Mathematically,

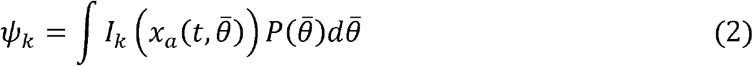

where *I_k_*(*x*) is an indicator function; *I_k_*(*x*) is equal to one if *x* lies in the *k*^th^ bin and zero otherwise.

The central idea behind MERIDIAN is to find the maximum entropy distribution 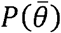 over parameters such that all predicted fractions *ψ_k_* agree with those estimated from experimental data, *ϕ_k_*. Formally, we seek 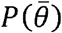 with the maximum entropy

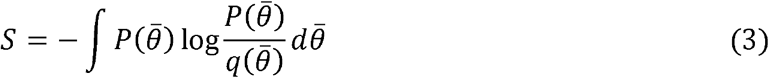

subject to normalization 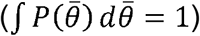 and data-derived constraints *ψ_k_* = *ϕ_k_* for all *k*. Here, 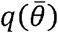 plays a role similar to the prior distribution in Bayesian approaches (Caticha and Preuss 2004). In this work, we choose 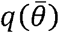 to be a uniform distribution within literature-derived ranges of parameters, but other choices can be implemented as well.

To impose aforementioned constraints and perform the entropy maximization, we use the method of Lagrange multipliers. To that end, we write the Lagrangian function

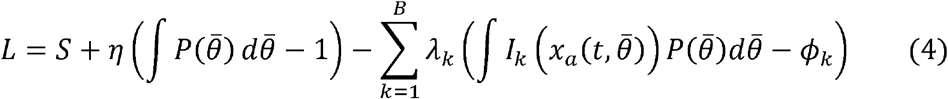

where *η* is the Lagrange multiplier associated with normalization *λ_k_* and are the Lagrange multipliers associated with fixing the predicted fractions *ψ_k_* to their experimentally measured values *ϕ_k_* in all bins. Differentiating equation (4) with respect to 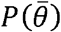 and setting the derivative to zero, we obtain

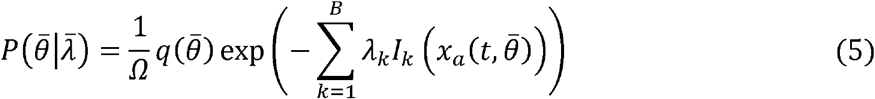

where *Ω* is the partition function that normalizes the probability distribution. Equation (5) is a key conceptual foundation of this work. We use it to estimate the parameter distribution from user-specified constraints. Here, for the sake of notational simplicity, we have restricted our discussion using measurement at a single time point. Later, we discuss how to generalize the approach when abundances of multiple species are measured at several time points (Methods equations (S24)).

Given the high dimensional nature of the parameter space in models of biological systems, the collected data is usually not sufficient to fully constrain the multidimensional parameter distribution (Banks 2012). As a result, the distribution inferred by MERIDIAN reflects both the true biological variability in parameters as well as parameter non-identifiability.

### Numerical estimation of Lagrange multipliers

The Lagrange multipliers in equation (5) need to be numerically optimized such that the predicted bin fractions are consistent with the experimentally estimated ones. Notably, the search for the Lagrange multipliers is a convex optimization problem (Methods) and we solve it using an iterative algorithm proposed in (Tkacik, Schneidman et al. 2006) (see Figure 2). Briefly, we start from a randomly chosen point in the space of Lagrange multipliers. In the *n*^th^ iteration of the optimization algorithm, using the current vector of the Lagrange multipliers 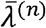, we estimate the predicted bin reactions using MCMC (Methods). Next, we estimate the error vector 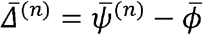 for the *n*^th^ iteration. We then update the multipliers for the *n*+1^st^ iteration as 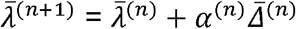 (see Figure 2). The positive “learning rate” *α*^(*n*)^ is chosen to minimize the error 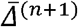 (Methods). Given that the data is likely to have experimental errors (see below) and that the signaling network model (described by equations (1)) is an approximation to a real biological system, it may not be possible to achieve a complete agreement between data and model fits. Thus, we terminate the iterative optimization procedure once the error reaches a predefined accuracy cutoff.

**Figure 2.**
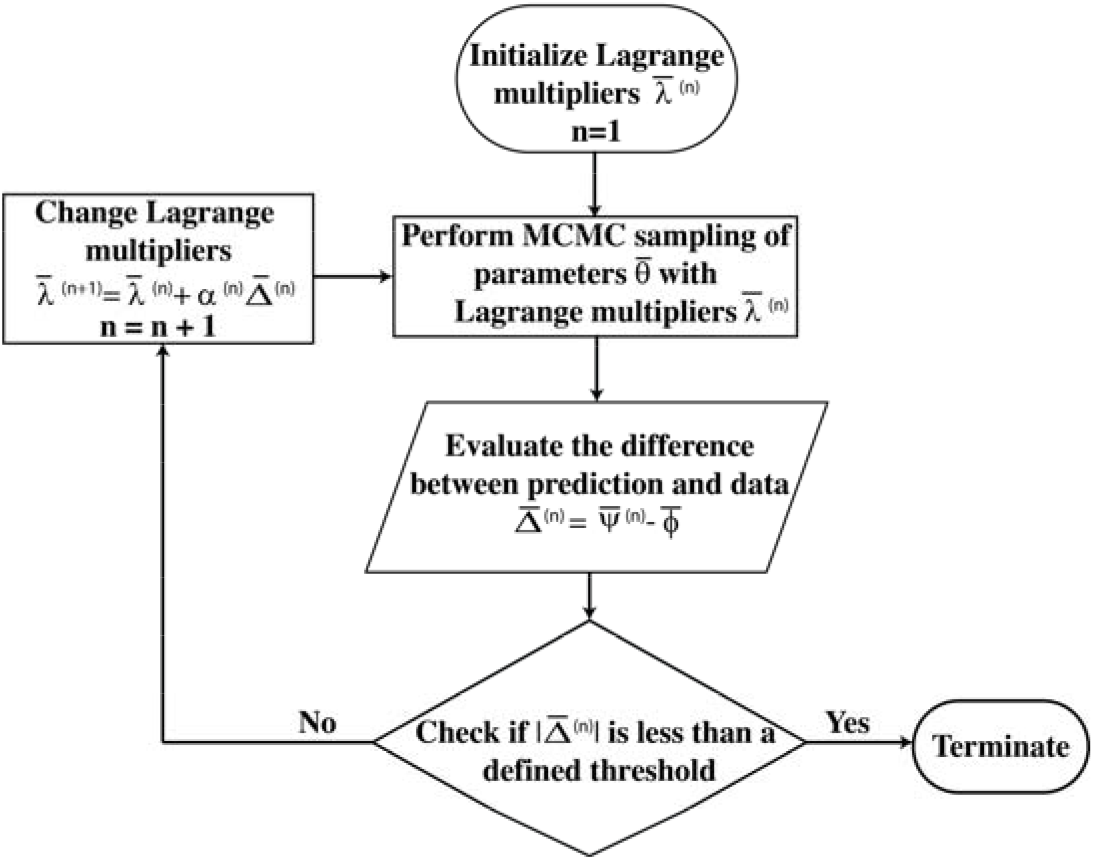
The workflow to numerically determine the values of Lagrange multipliers. In each iteration, we evaluate the error vector 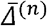 between the predicted bin fractions 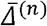 and the experimentally measured bin fractions using Markov chain Monte Carlo. We propose a new set of Lagrange multipliers based on the error vector. We repeat until the error reaches below a predefined accuracy cutoff.

**Figure 3.**
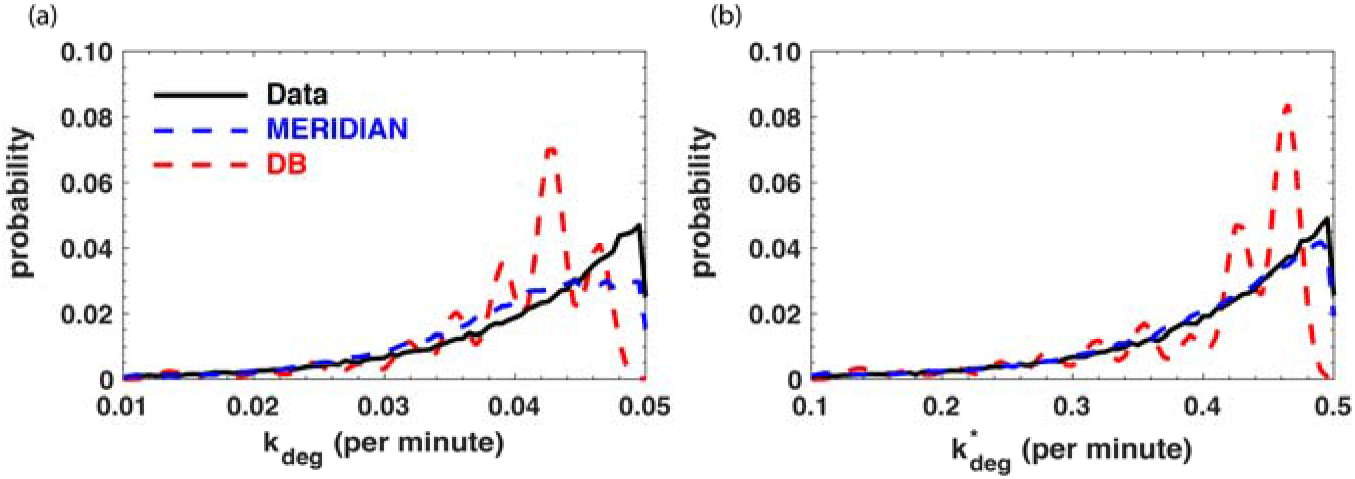
A comparison between the inferred parameter distributions and the data. We show the comparison between the true parameter distributions in panel a) and in panel b) (black lines) and the corresponding MERIDIAN-inferred distribution (blue lines) and the DB inferred distribution (dashed red lines).

### Making predictions using 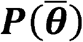

Here, we show how to make predictions using the inferred parameter distribution. In the discussion so far, we assumed that experimentally measured cell-to-cell variability had no errors. However, single cell experiments are often subject to uncertainty. Thus, we consider that the measurements are characterized by their mean values 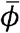 as well as the standard errors of the mean 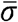, which are estimated using several experimental replicates.

Let us denote by 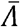 the vector of optimized Lagrange multipliers (see Figure 2), and by 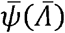 the predicted bin fractions of species variability across cells (Figure 1). We represent the experimental errors in the estimated bin fractions using a distribution over the Lagrange multipliers themselves. We express the posterior distribution over of Lagrange multipliers 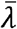 as (Methods)

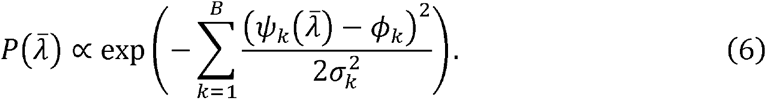

In equation (6), we have assumed (1) that the errors are normally distributed and (2) a uniform prior over Lagrange multipliers; both these assumptions can in principle be relaxed. The distribution over model parameters taking into account the errors in the measurement is then given by

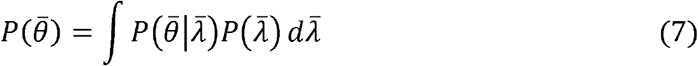

where 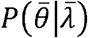 is given by equation (5) and the full distribution 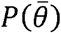 is given by averaging over the distribution of Lagrange multipliers 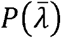.

Equations (5–7) can be used, in principle, to make model predictions. For example, consider any quantity of interest 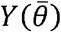 that depends on model parameters. The mean *m* of 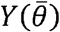 is then given by

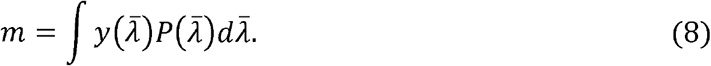

where

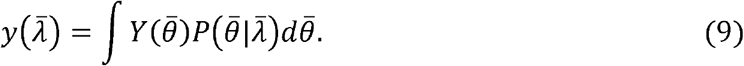

Similarly, the uncertainty *s* given by

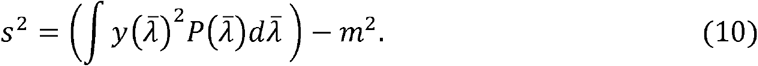

In practice it may be challenging to numerically integrate equations (8–10), because this requires sampling over multiple sets of Lagrange multipliers 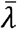, which in turn requires the estimation of 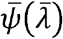 using MCMC (see equation (6)). However, estimates of the mean and the uncertainty can be analytically obtained if experimental errors are small (median error is 9% in our data). In this case, the model prediction *m* is simply given by:

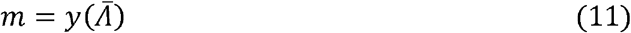

In equation (11), 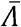 are the optimized Lagrange multipliers. Equation (11) is the maximum posterior estimate of equation (6). In Methods, we show that the estimate of the uncertainty *s* in the mean prediction is given by

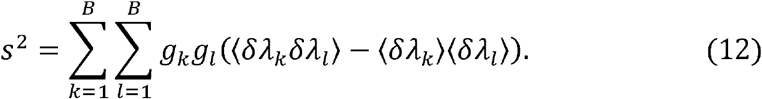

In equations (12) *δλ_k_* = *λ_k_* − *Λ_k_* and 〈*δλ_k_δλ_l_*〉 − 〈*δλ_k_*〉〈*δλ_l_*〉 is the corresponding covariance matrix among Lagrange multipliers. In equation (12), the “couplings” *g_k_* are given by

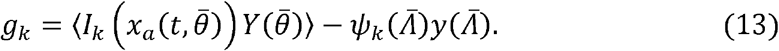

*g_k_* quantifies the correlation between the quantity of interest 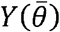 and the constraints 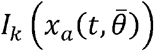. In Methods we show that

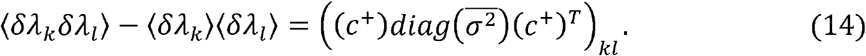

In equation (14) *c*^+^ is the Moore-Penrose pseudo-inverse of the covariance matrix *c. c* is given by:

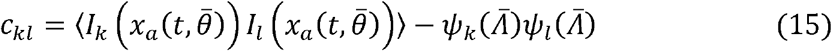

In equations (12–15), the averages denoted by angular brackets are calculated using equation (5) with the Lagrange multipliers 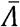 fixed at their optimized values.

To use the aforementioned approach in our calculations, we first estimate mean bin fractions 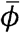 and the corresponding standard errors in the mean 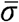 from experimental data. Then, by using the iterative scheme described in Figure 2, we estimate the optimized Lagrange multipliers 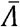 in equation (5) and the corresponding predictions 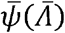. We then compute the covariance matrix *c* and its pseudo-inverse *c*^+^ (equation 15), and the covariance among the Lagrange multipliers (equation 14). Finally, for any quantity of interest 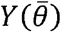, we compute the couplings 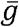 (equation 13). The predicted mean value and the associated uncertainty are given by equation (11) and equation (12) respectively.

### MERIDIAN performance on synthetic data

First, we use synthetic data to illustrate the utility of MERIDIAN and compare its performance with a previously developed discretized Bayesian (DB) approach by Hasenauer et al. (Hasenauer, Waldherr et al. 2011). In DB, a crucial step is to approximate the joint parameter distribution as a linear combination

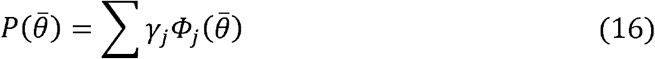

where 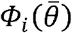 are a set of predefined distribution functions and *γ_i_* > 0 are the corresponding weights. In principle, if a sufficiently large number of such basis functions are chosen, their linear combination can capture any complex probability distribution. DB discretizes the multidimensional parameter space using a Cartesian grid and assumes that 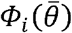 are multivariate Gaussian distributions centered at grid points. The weights *γ_i_* are sampled from their posterior distribution given the data. Because several other methods (Hasenauer, Waldherr et al. 2011, Hasenauer, Hasenauer et al. 2014, Loos, Moeller et al. 2018) also use linear combination of known functions to approximate the parameter distributions, a comparison with DB highlights the potential advantages of MERIDIAN.

To compare MERIDIAN with DB, we used a simplified growth factor network model (Methods, Supplemental Information Figure 1). Specifically, the network included three chemical species: the ligand *L*, ligand-free inactive receptors *R*, and ligand bound active receptors *P*. Ligand binding to receptors leads to their activation. Inactive receptors are constantly delivered to the cell surface, and active and inactive receptors are removed from the cell surface at different rates (Methods). In this model, the dynamics of the species was described by five parameters.

Using the aforementioned model, we generated synthetic single cell data by varying two key parameters: (1) rate of degradation of inactive receptors *k_deg_* and (2) rate of degradation of active receptors 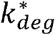. Notably, it was demonstrated that the processes that govern receptor trafficking on the cell surface are correlated at the single cell level (Kallenberger, Unger et al. 2017). To account for these correlations, we sampled the two parameters from a correlated bivariate exponential distribution (Iyer 2002). Using the joint parameter distribution we generated distribution of activated receptor levels corresponding to four different experimental conditions (ligand *L* = 2 ng/ml, *L* = 10 ng/ml, t = 10 minutes and steady state, Methods). Normally distributed random errors were then added to individual cell measurements. Mean bin fractions and standard errors in bin fractions were obtained using 5 separate synthetic datasets. These data were then used to infer the joint distribution 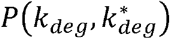 using both MERIDIAN and DB.

There are two types of hyper-parameters in DB: (1) number of Cartesian grid points and (2) the width of the multivariate Gaussian distributions. We performed DB-inference using several sets of these hyper-parameters. For each set, we first found the maximum likelihood weights *Γ_j_*. Next, we determined the optimal hyper-parameters using the Akaike information criterion (Akaike 1973). Using the optimal hyper-parameters, we sampled the posterior distribution over the weights *γ_j_* and estimated the joint distribution over parameters 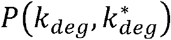 (Methods).

Notably, both MERIDIAN and DB were able to accurately fit the synthetic data (Methods, Supplemental Information Figure 2). The two approaches were also similar in capturing the Pearson correlation between the two rates (*ρ_DB_* ~ 0.19, *ρ_MERIDIAN_* ~ 0.18, and *ρ_Data_* ~ 0.3. However, they predicted substantially different parameter distributions. While MERIDIAN accurately captured both the distributions *P*(*k_deg_*) and 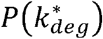, the distribution inferred using DB were very different from exponential distributions (Figure 4).

**Figure 4.**
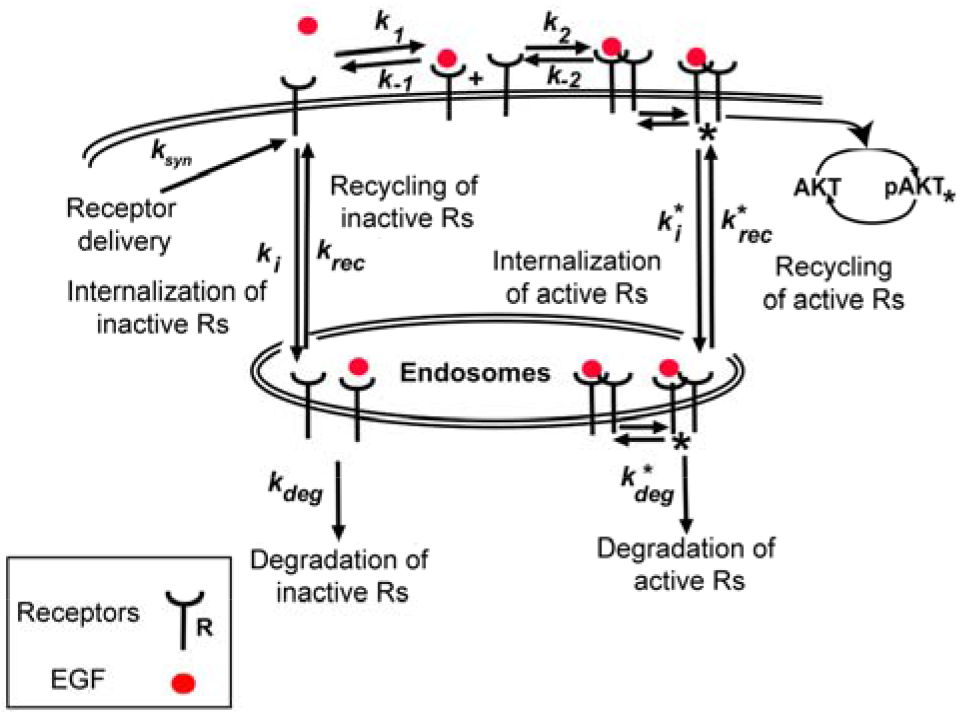
A schematic of the EGF/EGFR pathway leading to phosphorylation of Akt. Extracellular EGF binds to cell surface EGFRs leading to their dimerization. Dimerized EGFRs are autophosphorylated and in turn lead to phosphorylation of Akt. Receptors are also removed from cell surface through internalization into endosomes. See Methods for details of the model.

Next, we obtained the inferred parameter distributions using DB for several sets of inference hyper-parameters by sampling over the posterior distribution over the weights (Supplemental Information Table 4). We quantified the similarity between the inferred and true distributions using three different cumulative-based metrics. Notably, MERIDIAN outperformed a vast majority of the tested choices of DB hyper-parameters (more than 93%) (Methods, Supplemental Information Table 4, and Supplemental Information Figure 3).

Finally, we note that applying the DB approach to realistic models of biological networks (with 10-30 parameters) may be computationally prohibitive. If we choose *N_G_* grid points per dimension in an inference of a *K* dimensional parameter space, DB employs 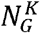 basis functions. For example, for a signaling network described by 20 parameters, employing 5 grid points per dimension will require ~10^14^ basis functions.

### Using MERIDIAN to study EGFR/Akt signaling

#### Computational model of the EGFR/Akt signaling network

Signal transduction in the EGFR/Akt network is illustrated in Figure 4. Following stimulation of cells with EGF, the ligand binds to the cell surface EGFRs. Ligand-bound receptors dimerize with other ligand-bound receptors as well as ligand-free receptors. EGFR dimers then phosphorylate each other and phosphorylated receptors (active receptors, pEGFRs) on the cell surface lead to downstream phosphorylation of Akt (pAkt). Both active and unphosphorylated (inactive) receptors are internalized with different rates from the cell surface through receptor endocytosis. After addition of EGF in the extracellular medium, pAkt levels increase transiently within minutes and then, as a result of receptor endocytosis and action of phosphatases, both pAkt and surface EGFR (sEGFR) levels decrease within hours after EGF stimulation (Chen, Schoeberl et al. 2009).

To explore the cell-to-cell variability in this pathway, we used a dynamical model of EGF/EGFR dependent Akt phosphorylation based on Chen et al. (Chen, Schoeberl et al. 2009). The model includes reactions describing EGF binding to EGFR and subsequent dimerization, phosphorylation, dephosphorylation, internalization, and degradation of receptors. To keep the model relatively small, we simplified pEGFR-dependent phosphorylation of Akt by assuming a single step activation of Akt by pEGFR (Methods). We note that the first and second order rate constants employed the model should be treated as effective rates given that the law of mass action is only an approximation to describe the complex processes in the EGFR/Akt pathway. The model had 17 chemical species and 20 parameters. See Supplemental Information Table 2 for a list of model parameters and Supplemental Information Table 3 for a list of model variables. The model equations are given in Methods.

#### Numerical inference of the parameter distribution

To estimate the signaling parameter distribution consistent with experimental data using MERIDIAN, we used experimentally measured cell-to-cell variability in pAkt levels at early times after EGF stimulation. Specifically, we used measured pAkt levels after stimulation with five different EGF doses (0.1, 0.316, 3.16, 10, and 100 ng/ml) at 4 early time points (5, 15, 30, and 45 minutes) (Lyashenko, Niepel et al. 2017). Additionally, we used sEGFR levels without EGF stimulation and after 3 hours of EGF stimulation at 1 ng/ml. We used 11 bins to represent each experimentally measured distribution; the bin sizes and locations were chosen to cover the entire range of observed variability (Supplemental Information Table 1). There were a total of 264 bin fractions and corresponding 264 Lagrange multipliers. We numerically determined the optimal Lagrange multipliers corresponding to the bin fractions using the procedure described above (see Figure 2, Methods). It took approximately 90 hours to learn the optimal Lagrange multipliers.

Notably, the optimal Lagrange multipliers accurately reproduced the experimentally measured bin fractions (Pearson’s *r*^2^ = 0.9, *p* < 10^−10^, median relative error ~ 14%). Furthermore, fitted bin fractions obtained in two independent calculations showed excellent agreement with each other as expected for a convex optimization problem (Pearson *r*^2^ = 0.99, *p* < 10^−10^, Supplemental Information Figure 4). In Figure 5, we show the temporal profile of measured cell-to-cell variability in pAkt levels (colored circles) at EGF stimulation of 10 ng/ml and the corresponding fits (dashed black lines) based on the inferred parameter distribution. The fits to all 24 distributions are given in Supplemental Information Figure 5. The marginal distributions of the individual parameters are given in Supplemental Information Figure 6 and the correlation structure amongst parameters is given in Supplemental Information Table 7.

**Figure 5.**
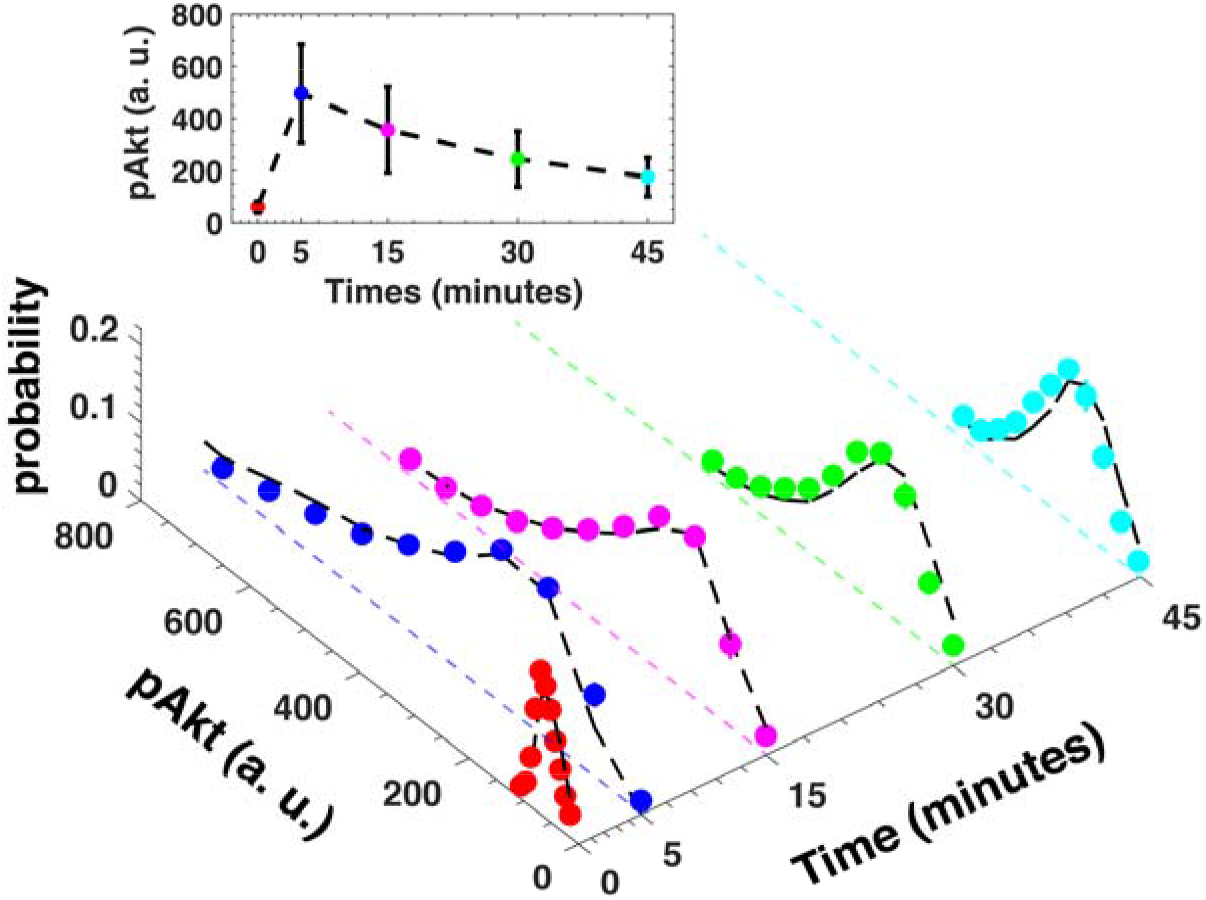
Experimental estimated cell-to-cell variability in pAkt levels used to infer the parameter distribution. We show distribution of pAkt levels at 0, 5, 15, 30, and 45 minutes after exposure to 10 ng/ml EGF. The colored circles represent the experimentally measured pAkt distributions used in the inference of the parameter distribution. The black dashed lines represent fitted distributions. The inset shows the experimentally measured population average pAkt levels at multiple time points. Error bars represent standard deviation. Error bars in the inset represent population standard deviations.

#### Prediction of single cell dynamics

Akt is a key hub of mammalian cell signaling (Manning and Toker 2017). Naturally, sustained activity of phosphorylated Akt (pAkt) is implicated in diverse human diseases, such as psychiatric disorders (Gilman, Chang et al. 2012) and cancer (Vivanco and Sawyers 2002). Using the developed approach, we investigated whether we could predict pAkt levels hours after EGF stimulation using the parameter distribution inferred from pAkt variability at early times after EGF stimulation. To that end, we numerically sampled multiple parameter sets using the inferred parameter distribution and predicted pAkt levels at late time across a range of EGF stimulation levels corresponding to each parameter set. We compared predicted and experimentally observed distribution of pAkt levels across cells at late times (180 minutes) after sustained EGF stimulation (Figure 6a,b, Supplemental Information Figure 7). Our simulations correctly predicted that a significant fraction of cells have high pAkt levels hours after stimulation; the predicted and observed coefficient of variation (CV) of the pAkt distributions in cells stimulated with 10 ng/ml EGF for 180 minutes were in good agreement, 0.41 and 0.37 respectively. Notably, the inferred parameter distribution accurately captures the population mean and variability (Figure 6c) in pAkt levels at late times across four orders of magnitude of EGF concentrations used to stimulate cells.

**Figure 6.**
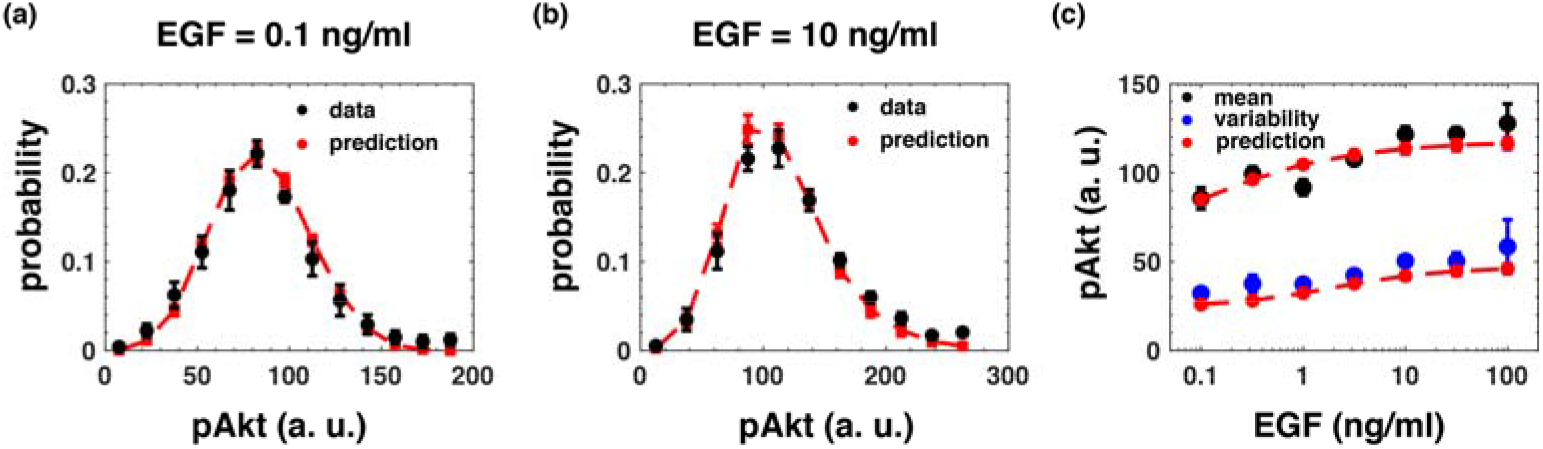
Prediction pAkt levels at late times. (a and b) Measured distributions (black circles and lines) and the corresponding predictions (red circles and dashed lines) of cell-to-cell variability in pAkt levels at 180 minutes after stimulation with 0.1 ng/ml and 10 ng/ml EGF respectively. (c) Measured mean pAkt levels (red circles) and measured standard deviation in pAkt levels (blue circles) at 180 minutes after sustained stimulation with EGF (x-axis) and the corresponding predictions (dashed red lines). The error bars in experimental data represent standard deviation. The error bars in model predictions represent the estimated uncertainty.

Importantly, MERIDIAN allowed us to investigate the biochemical parameters that significantly correlate with high pAkt levels at steady state. Interestingly, across all simulated trajectories, the levels of cell surface EGFR showed the highest correlation with pAkt levels among all receptor-related parameters (Supplemental Information Table 6, Pearson *r* = 0.4, EGF stimulation 10 ng/ml). This suggests that cells with high EGFR levels likely predominantly contribute to the sub-population of cells with high steady state pAkt activity. This demonstrates how MERIDIAN can be used to gain mechanistic insight into heterogeneity in signaling dynamics based on single cell data.

We next investigated whether MERIDIAN could predict the heterogeneity in EGFR levels after prolonged stimulation with EGF. To that end, we compared the predicted and the experimentally measured the steady state EGFR levels across EGF stimulation doses. Similar to pAkt, the simulations accurately captured both the population mean and variability of the EGFR receptor levels across multiple doses of EGF stimulations (Figure 7c). The simulations and experiments demonstrate that in agreement with model prediction that even hours after the growth factor stimulation there is a significant fraction of cells with relatively high levels of EGFR (Figure 7a,b).

**Figure 7.**
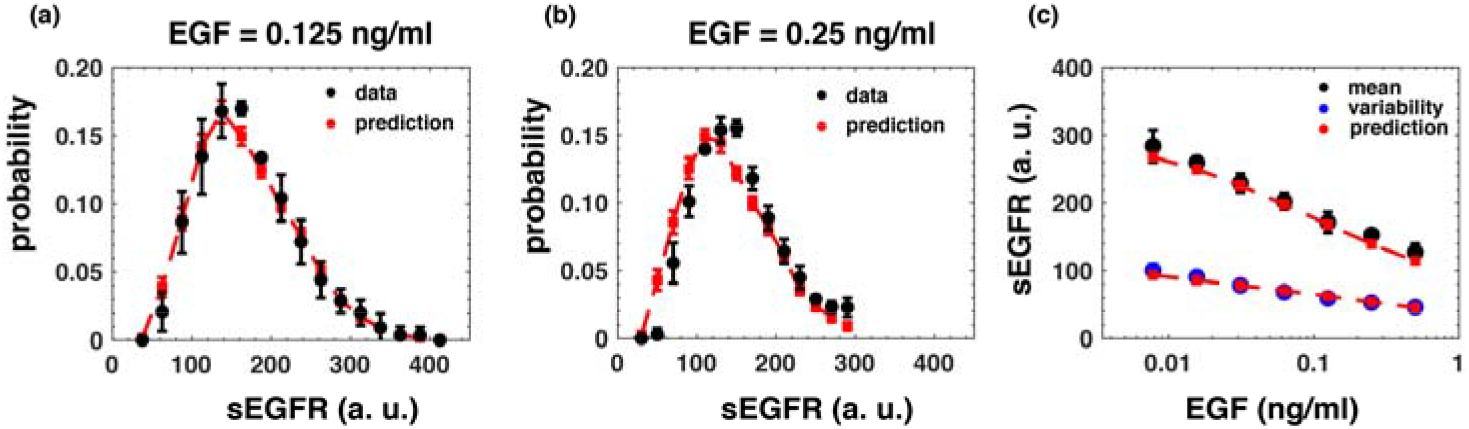
Prediction of sEGFR levels at late times. (a and b) Measured distributions (black circles and lines) and the corresponding predictions (red circles and dashed lines) of cell-to-cell variability in sEGFR levels at 180 minutes after stimulation with 0.125 ng/ml and 0.25 ng/ml EGF respectively. (c) Measured mean sEGFR levels (red circles) and measured standard deviation in sEGFR levels (blue circles) at 180 minutes after sustained stimulation with EGF and the corresponding predictions (dashed red lines). The error bars in experimental data represent standard deviation. The error bars in model predictions represent the estimated uncertainty.

### Possible extensions of the MERIDIAN framework

#### Using MERIDIAN with inherently stochastic networks

A straightforward extension makes it possible to use the MERIDIAN framework for signaling networks when the time evolution of species abundances is intrinsically stochastic, for example, transcriptional networks and prokaryotic signaling networks with relatively small species abundances (Raj and van Oudenaarden 2008). To that end, we can modify the definition of the predicted bin fraction 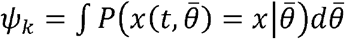 where 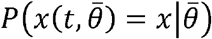 is the distribution of *x* values at time t with parameters 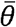. The distribution can be obtained numerically using Gillespie’s stochastic simulation algorithm (Gillespie 2007) and its fast approximations (Cao and Grima 2018) or approximated using moment closure techniques (Gillespie 2009). Notably, we have previously implemented this generalization of MERIDIAN to understand intrinsic and extrinsic noise in a simple gene expression circuit in *E. coli* (Dixit 2013).

#### Constraining moments in MERIDIAN

MERIDIAN can also be used to infer parameter distributions when, instead of the entire abundance distributions only a few moments of the distribution are available, such as average protein abundances measured using quantitative western blots or mass spectrometry (Shi, Niepel et al. 2016). For example, we consider the case where the population mean *m* and the variance *v* of one species *x* are measured at a fixed time point *t*. Instead of constraining fractions *ψ_k_* that represent cell-to-cell variability in different bins of the relevant abundance distribution, we can constrain the population mean 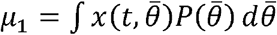 and the second moment 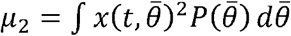 to their experimentally measured values *m* and *v+m*^2^ respectively. Entropy maximization can then be carried out with these constraints. In this case, we have

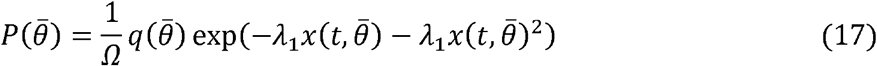

### Using MERIDIAN with high-dimensional data

MERIDIAN can be used to infer parameter distributions when multiple chemical species are experimentally measured in single cells at the same experimental condition. It may be difficult to accurately estimate the multidimensional bin counts from multidimensional data. Therefore, one can apply the following approach. For example, two species *x* and *y* simultaneously are measured across several cells, in addition to constraining the onedimensional bin fractions 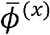 and 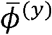, we can also constrain the cross-moment *r* = 〈*xy*〉. With these three types of constraints, the maximum entropy distribution is given by

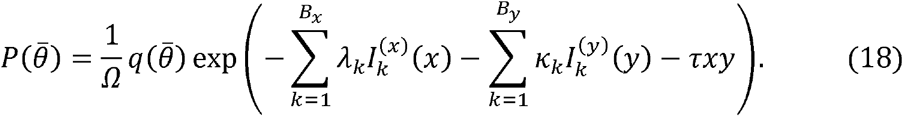

In equation (18), 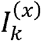 and 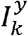 are the indicator functions for species *x* and *y* respectively, *B_x_* and *B_y_* are the number of bins used in the *x*- and the *y*-dimension, 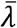 and 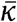 are Lagrange multipliers constraining the bin fractions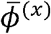 and 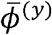 respectively, and *τ* is the Lagrange multiplier that constrains the cross-moment. By adding cross-moment constraints for each pair of species, equation (18) can be easily generalized to multiple dimensions; this will add ~*N*^2^/2 Lagrange multipliers where *N* is the number of measured species.

### Speeding up MERIDIAN inference using neural networks

A key numerical bottleneck in applying the MERIDIAN inference approach is the numerical optimization of a large number of Lagrange multipliers. It is a well-known problem in maximum entropy inference (Loaiza-Ganem, Gao et al. 2017). To address this problem, recently Loaiza-Ganem et al. (Loaiza-Ganem, Gao et al. 2017) proposed a maximum entropy flow network approach based on approximate deep generative modeling. Briefly, instead of finding the continuous density of the maximum entropy distribution in equation (5), they find the approximate maximum entropy distribution within a parametric family. The family is parameterized by several layers of a neural network and was shown to be sufficiently accurate in approximating true maximum entropy distributions. Moreover, a recent extension of this approach (Bittner and Cunningham 2019) enables fast simultaneous sampling of maximum entropy distributions along with a distribution of Lagrange multipliers (equation (7)). We believe that these fast methods will be crucial when using the MERIDIAN approach to study large signaling networks with several experimentally measured single cell distributions.

### Using MERIDIAN with live cell imaging data

Finally, we discuss how we can use MERIDIAN to infer parameters from experiments where dynamics of species abundances within single cells are measured using live cell imaging (Meyer, D’Alessandro et al. 2012, Kallenberger, Unger et al. 2017). For example, consider that the time evolution of a species *x*(*t*) is measured in *n_c_* cells from time *t* = 0 to *t* = *T*. We can discretize the continuous time observations into K discrete times {*t*_1_, *t*_2_,…, *t_k_*}. At each time point *t_i_*, we can then divide the range of observed abundances in *B_i_* bins. Then, each individual dynamical trajectory *x*(*t*) can be characterized by a vector of indices *x*(*t*) ~ {*B*_1*a*_1__, *B*_2*a*_2__,…, *B_ka_k__* where *B_ia_i__* is the index of the abundance distribution bin through which the trajectory *x*(*t*) passed at time point *t_i_*. Given a sufficiently large number of trajectories, we then can constrain the fraction of trajectories that populate a given sequence of bins to infer the parameter distribution.

## Discussion

Cells in a population exhibit heterogeneity in part because of heterogeneity in signaling network parameters (Albeck, Burke et al. 2008, Niepel, Spencer et al. 2009, Spencer, Gaudet et al. 2009, Meyer, D’Alessandro et al. 2012, Llamosi, Gonzalez-Vargas et al. 2016, Kallenberger, Unger et al. 2017). In this work, we developed a maximum entropy based approach to infer this parameter heterogeneity from single cell measurements of chemical species abundances. Notably, the inferred distribution combines two components: (1) the true biological parameter variability due to cell-to-cell heterogeneity and (2) the non-identifiability in parameter estimation given the single cell data. Consequently, the inferred distribution is likely to be broader compared to the true biological variability (Mukherjee, Seok et al. 2013). Notably, the non-identifiability component can be further minimized by (1) optimally designing experimental conditions to reduce non-identifiability (Bandara, Schloder et al. 2009, Kreutz and Timmer 2009) or by (2) directly including constraints on population average measurements of rate constants and other parameters of the signaling network.

We briefly discuss key differences between MERIDIAN and a previous maximum entropy based approach by Waldherr et al. (Waldherr, Hasenauer et al. 2009). Waldherr et al. employ the so-called Latin hypercube sampling (LHS) approach (Stein 1987). A potential advantage of LHS is that it avoids computationally expensive determination of the Lagrange multipliers. At the same time, LHS only sparsely samples the parameter space and generally cannot assign probabilities to arbitrary high dimensional parameter points. In contrast, an advantage of MERIDIAN is that the continuous density defined in equation (5) allows us to estimate the relative probability of any parameter point. Finally, unlike the LHS approach, MERIDIAN allows us to estimate the uncertainty in model predictions using measurement errors.

Recent developments in cytometry (Chattopadhyay, Gierahn et al. 2014) and single cell RNA sequencing (Saliba, Westermann et al. 2014) make it possible to simultaneously quantify multiple species abundances in single cells. Elegant statistical approaches have been developed to reconstruct trajectories of intracellular species dynamics consistent with time-stamped single cell abundance data (Gut, Tadmor et al. 2015, Mukherjee, Jensen et al. 2017, Mukherjee, Stewart et al. 2017). Complementary to these statistical methods, our approach (1) allows us to infer the distribution over signaling parameters that describe mechanistic interactions in the signaling network and moreover (2) the inferred parameter distribution can be used to predict the ensemble of single cell trajectories for time intervals and experimental conditions beyond the measured abundance distributions.

In this work, we applied the developed framework to signaling network data. However, it can also be used in other diverse research contexts. For example, the framework can be applied to computationally reconstruct the distribution of longitudinal behaviors from cross-sectional time-snapshot data in fields such as public health, economics, and ecology or to estimate parameter distributions from lower dimensional statistics (Das, Mukherjee et al. 2015).

## Methods

### Experimental Details

In this work, we used distributions of cell-to-cell variability in phosphorylated Akt levels as well as cell surface EGFR levels. We used the experimental data on pAkt levels previously measured in Lyashenko *et al*. (Lyashenko, Niepel et al. 2017) We measured cell-to-cell variability in sEGFR levels in this work. Briefly, we describe the experimental methods here.

MCF 10A cells (Soule, Maloney et al. 1990) were obtained from the ATCC. The cells were grown according to ATCC recommendations. We confirmed the cell identity by short tandem repeat (STR) profiling at the Dana-Farber Cancer Institute. We tested the cells with MycoAlert PKUS mycoplasma detection kit (Lonza) and ensured that they were free of mycoplasma infection. For the experiments, we coated 96 well plates (Thermo Fisher Scientific) with type I collagen from rat tail (Sigma-Aldrich) by incubating plates with 65 microliter of 4mg/ml collagen I solution in PBS for two hours at room temperature. We washed the plates twice with PBS using EL406 Microplate Washer Dispenser (BioTek) and sterilized them under UV light for 20 minutes prior to use. Cells were harvested during logarithmic growth. We dispensed 2500 cells per well into collagen-coated 96 well plates using a EL406 Microplate Washer Dispenser. We grew the cells in 200 microliter of complete medium for 24 hours. The cells were serum-starved twice in starvation media (DMEM/F12 lemented with 1% penicillin-streptomycin and 0.1% bovine serum albumin). Next, we incubated the cells in 200 microliter of starvation media for 19 hours and again for one more hour. This time point constituted t=0 for all experiments.

We created the EGF treatment solutions by dispensing the appropriate amounts of epidermal growth factor (EGF, Peprotech) into starvation media using a D300 Digital Dispenser (Hewlett-Packard). To fit the parameter distributions, we used EGF concentrations of 0.1, 0.31, 3.16, 10, and 100 ng/ml for Akt phosphorylation measurements and EGF concentrations of 0, 1, and 100 ng/ml for surface EGFR measurements. At t=0 cells were stimulated with 100 microliter of 3x solution and incubated for indicated times (5, 15, 30, and 45 minutes for pAkt and 180 minutes for sEGFR). To test the model predictions, we collected pAkt distributions at 90 and 180 minutes after stimulation with 0.01, 0.031, 0.1, 0.31, 1, 3.16, 10, 31.6, and 100 ng/ml EGF. We also measured sEGFR distributions at 180 minutes after stimulation with 0.0078, 0.0156, 0.0312, 0.0625, 0.125, 0.25, 0.5, 1, and 100 ng/ml of EGF. All incubations were terminated by adding 100 μl of 12% formaldehyde solution (Sigma) in phosphate buffered saline (PBS) and fixing the cells for 30 min at room temperature.

We performed all subsequent washes and treatments with the EL406 Microplate Washer Dispenser. We washed the cells twice in PBS and permeabilized them with 0.3% Triton X-100 (Sigma-Aldrich) in PBS for 30 min at room temperature. Cells were washed once again in PBS, and blocked in 40 microliter of Odyssey blocking buffer (LI-COR Biotechnology) for 60 min at room temperature. Cells were incubated with 30 microliter of anti-phospho-Akt (Cell Signaling Technologies, #4060, 1:400) or anti-EGFR (Thermo Fisher Scientific, MA5-13319, 1:100) over night at 4°C. We then washed the cells once in PBS and three time in PBS with 0.1% Tween 20 (Sigma-Aldrich; PBS-T for 5 min each and incubated with 30 microliter of a 1:1000 dilution of Alexa Fluor 647 conjugated goat anti-rabbit or goat anti-mouse secondary antibody in Odyssey blocking buffer for 60 min at room temperature. Next we washed the cells two times in PBS-T, once with PBS, and stained for 30 min at room temperature with whole cell stain green (Thermo Fisher Scientific) and Hoechst (Thermo Fisher Scientific). Finally, cells were washed three times in PBS, covered in 200 microliter of PBS, and sealed for microscopy. We imaged cells with an Operetta high content imaging system (Perkin Elmer) and analyzed the resulting scans using the Columbus image data storage and analysis system (Perkin Elmer). We performed the experiments in biological triplicates for surface EGFR and quadruplets for pAkt. To avoid potentially pathological bright cells, we removed the top 1% of the data in all single cell distributions.

### Computational details

#### Incorporating experimental error in MERIDIAN

In this section, we expand on the mathematical details of how to incorporate experimental errors in the MERIDIAN inference procedure. As in the main text, we denote by 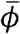 the experimentally estimated mean bin fractions and by 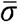 the corresponding standard errors in the mean. We assume that following an iterative procedure described in Figure 2, we have obtained an optimal set of Lagrange multipliers 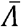. We denote by 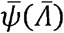 the corresponding model predicted bin fractions.

Any fixed set of Lagrange multipliers uniquely determines model predictions 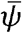. Thus, the errors in experimental measurements are captured by a distribution over the Lagrange multipliers themselves. We write the probability of non-optimal Lagrange multipliers 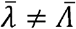 as

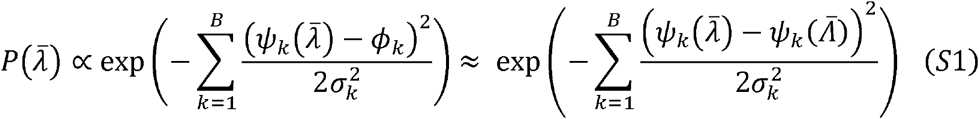

Equation (S1) assumes that the errors are normally distributed and that the residuals 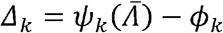 are small. We have also neglected the Jacobian determinant associated with changing the variables from 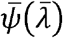 to 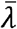. Sampling Lagrange multipliers from equation (S1) is in principle possible but may be numerically inefficient. This is because it requires on-the-fly estimation of predicted bin fractions 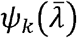 for non-optimal Lagrange multipliers 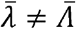. However, if we are interested the first two moments (means and uncertainties), we can approximate the distribution over Lagrange multipliers as a multivariate Gaussian distribution. This is equivalent to assuming that the experimental errors *σ_k_* are small compared to the mean values *ϕ_k_*. In the EGFR/Akt data used in this work, the standard errors in the mean are indeed small; median relative error is ~9% and the mean relative error is ~11%. To express the distribution in equation (S5) as a Gaussian, we first write

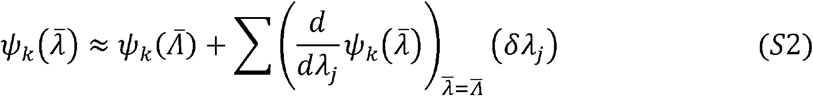

where *δλ_j_* = *λ_j_* − *Λ_j_* is the deviation in *λ_j_* away from the optimal Lagrange multipliers. Using linear response theory from statistical physics (Hazoglou, Walther et al. 2015), the derivatives in equation (S2) can be expressed as ensemble average over the parameter space. We write

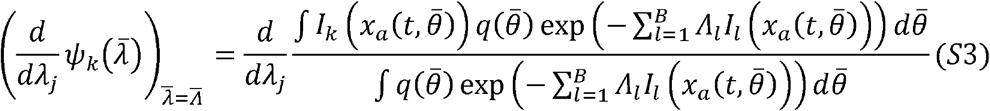

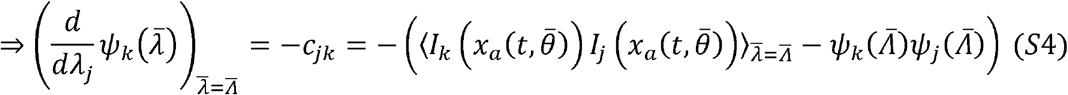

In equation (S4), *c_jk_* is the covariance matrix among the constraints. The average is computed using equation (5) in the main text with 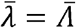.

Combining equations (S1), (S2), and (S4), we obtain the Gaussian approximation to the distribution over Lagrange multipliers,

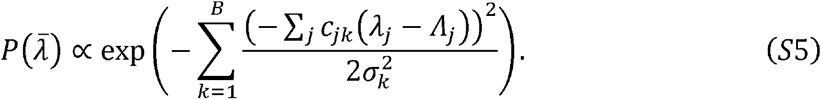

The multivariate Gaussian distribution in equation (S5) is fully determined by the means and the covariance matrix of the Lagrange multipliers. We determine these next.

Since we assume that the model can fit the data reasonably accurately, the average value of the deviation in Lagrange multipliers in equation (S5) is 〈*δλ_j_*〉 = 0. Next, we estimate the covariance matrix among the Lagrange multipliers. Let us consider a particular bin fraction *ϕ_k_*. The model estimated uncertainty is given by

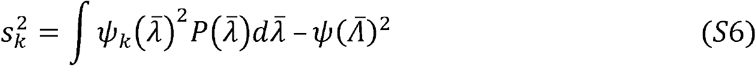

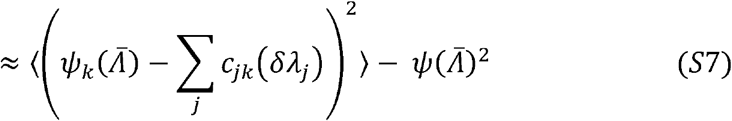

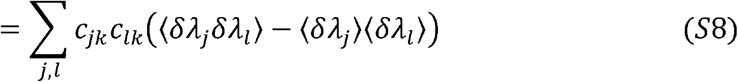

The experimentally estimated uncertainty in *ϕ_k_* is 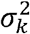. Equating the two, we have

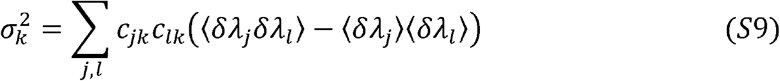

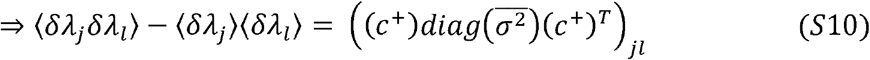

In equation (S10), *c*^+^ is the pseudoinverse of the covariance matrix.

These first two moments fully describe the multivariate Gaussian distribution over Lagrange multipliers (equation (S5)). Next, we show how to estimate mean predictions and uncertainty in model predictions.

Consider a variable 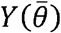 that depends on model parameters 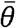. We are interested in estimating its mean predicted value “m” and the corresponding uncertainty “s”. Let us denote by 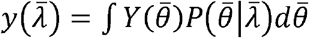 the model prediction when the Lagrange multipliers are fixed at 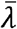. We have the mean prediction

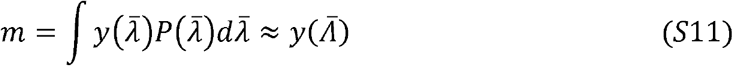

Next, we seek the estimated uncertainty (see equation 10 in the main text),

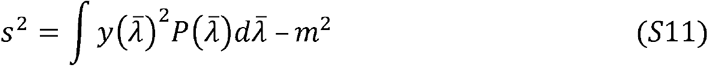

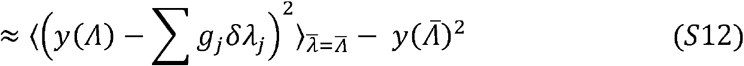

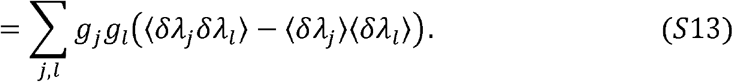

In equation (S13), the couplings *g_j_* are given by

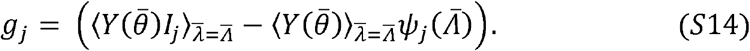

Equations (S11–S14)) show how to estimate model predictions and the corresponding uncertainty from the parameter distribution 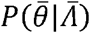 (equation (5) in the main text).

In the theoretical development above, we restricted the Taylor series expansion to the first order in 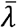. More generally, higher order Taylor series expansions can also be included. Notably, similar to equation (S4), all higher order Taylor series coefficients can be estimated using MCMC calculations performed using the parameter distribution 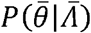.

### Comparison between DB and MERIDIAN

#### Simplified signaling network

In this section, we describe the details of the simple growth factor model used to generate synthetic data and the procedure to fit data to infer the parameter distribution using the maximum entropy approach as well as the discretized Bayesian approach of Hasenauer et al. (Hasenauer, Waldherr et al. 2011).

The model comprised three species: the extracellular ligand *L*, inactive ligand-free cell surface receptors *R*, and active ligand-bound cell surface receptors *P*. The dynamics of the three variables was dictated by five parameters: the rate of ligand binding to inactive receptors *k*_1_, the rate of ligand unbinding from active receptors *k*_−1_, the rate of removal of inactive receptors *k_deg_*, the rate of removal of active receptors 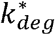, and the steady state cell surface receptor level in the absence of the ligand *R_T_*. See Supplemental Information Figure 1 for an illustration.

In the model, the dynamics of the species are governed by the following two equations:

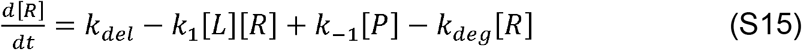

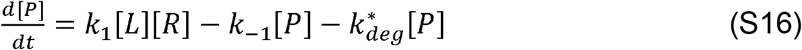

We simulated the model to generate synthetic data as follows. We fixed three parameters 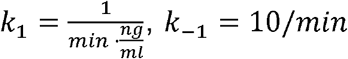, and *R_T_* = 5. We generated single cell distribution by varying the rates *k_deg_* and 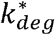. The rate of delivery of receptors to the cell surface was set equal to *k_del_* = *R_T_k_deg_*. The rates were sampled from a correlated bivariate exponential distribution (Iyer 2002). *k_deg_* was restricted between 0.01/min and 0.05/min and 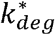 was restricted between 0.1/min and 0.5/min. The Pearson correlation between the sampled rates was set to be 0.28. Using 50000 sampled rates, we solved the differential equations (S15) and (S16) and generated single cell data on the number of activated receptors at four conditions (*L* = 2 ng/ml and 10 ng/ml, and *t* = 10 minutes, and steady state). A normally distributed random error was added to each single cell readout *P* that represented 5% of the magnitude of the data point (*P* → *P*\* (1 + 0.05*X*) where *X* was a standard normal variable). The 50000 single cells were split into five experiments of 10000 cells each. From these five experiments, we estimated the mean bin fractions 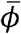 and the corresponding standard errors of the means 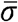. There were a total of 44 bin fractions (Supplemental Information Table 4).

We used these bin fractions to infer the parameter distribution using MERIDIAN and a previously developed discretized Bayesian (DB) approach.

#### Implementation of DB

In the DB approach, we approximated the parameter distribution as a linear combination

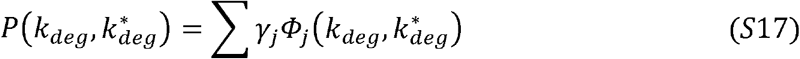

where 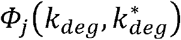 are bivariate Gaussian distributions situated on an equally spaced *N*×*N* Cartesian grid and *γ_j_* are the corresponding weights. Here, *N* is a hyper-parameter of the inference procedure. In order to test the accuracy of the DB approach, we used multiple values of the number of grid points (*N* = 5, 10, 15, and 20). Another choice of hyper-parameter is the covariance matrix of the Gaussians. We tested the covariance matrix of the Gaussians as: 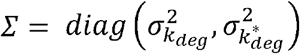 where 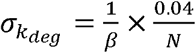 and 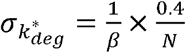. The hyper-parameter *β* controls the width of the bivariate Gaussian distribution. We scanned over multiple values of the parameter *β* (*β* = 1, 2, 4, 8, 16, and 32). Overall, we performed 24 DB inference calculations (4 N values × 6 *β* values).

In any given calculation, for every value of the index *j* (indicating one Gaussian basis function) we sampled *N_sim_* = 10^4^ parameter points from the bivariate Gaussian distribution 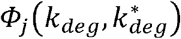 and estimated the predicted bin fractions 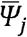. For any set of weights {*γ_j_*} the overall predicted bin fractions for the *i*^th^ bin is then expressed as the linear combination

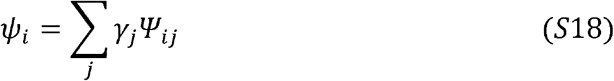

In order to decide which DB hyper-parameters to use for further analysis, we used the Akaike information criterion (AIC) (Akaike 1973). The AIC balances the accuracy with which a model fits the data and the number of parameters in the model. We determined the AIC for a given set of DB hyper-parameters as follows. First, we found the maximum likelihood set of weights by maximizing the log-likelihood

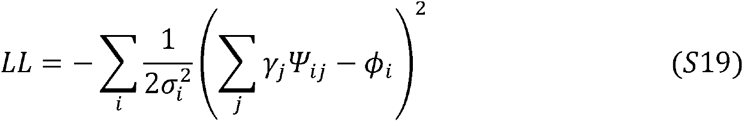

In equation (S19), *ϕ_i_* is the mean “experimental” bin fractions in the *i*^th^ experiment and is the corresponding standard error of the mean. Next, we calculated the AIC as AIC = 2*N*^2^ − 2*LL* where *N^2^* was the total number of free parameters in each model (the *N*^2^ weights). For further analysis, we used the set of hyper-parameters that minimized the AIC. This corresponded to *N* = 10 grid points per dimension and *β* = 4. The statistical characterization of all tested models is presented in Supplemental Information Table 4.

To account for the non-identifiability in the parameter distribution, we sampled the full posterior distribution over the weights using MCMC. The posterior distribution was given by the exponential of the log likelihood in equation (S19). For this calculation, we used a total of 4×10^7^ MCMC steps in the space of weights *γ_j_*. The first half of the simulation was discarded as equilibration. In the second half of the MCMC, we stored weights every 10000^th^ step. For each set of weights, we sampled 250 bivariate Gaussian variables by first sampling 50 grid points according to a multinomial distribution with the weights as probabilities and then sampling 5 bivariate Gaussian variables per sampled grid point.

#### Implementation of MERIDIAN

The optimal Lagrange multipliers 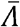 for MERIDIAN were inferred using an iterative procedure described below. In the iterative search, the predicted bin fractions 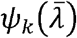 were estimated using direct integration using the trapezoidal rule. We note that integration using trapezoidal rule is not possible for a high dimensional parameter space. In order to incorporate the errors in experimental data, we sampled over the posterior distribution over Lagrange multipliers. First, we approximated the posterior distribution as a multivariate Gaussian distribution (see equation (S5), (S9), and (S14)). Next, we sampled 5000 sets of Lagrange multipliers using this multivariate Gaussian distribution. For each set of Lagrange multipliers, we performed MCMC calculations using the Metropolis criterion (see below for details) to samples through the parameter space using equation (5) in the main text. Briefly, 10^5^ steps of MCMC were used for equilibration and 10^4^ steps were used to save parameters. Parameters were saved every 500^th^ step.

#### Comparing MERIDIAN and DB

We used three different metrics to compare the marginal parameter distributions *P*(*k_deg_*) and 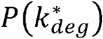 predicted using MERIDIAN and DB. First, we collected 50000 parameter samples from the parameter distributions using both MERIDIAN and DB. Next, we binned these samples using intervals 0.01:0.0005:0.05 and 0.1:0.005:0.5 for *k_deg_* and 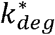 respectively using the hist function in MATLAB. Using these bins, we found the cumulative distributions *F_M_*(*k_deg_*) and 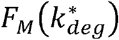 respectively. Here, the subscript “M” denotes the method (MERIDIAN and DB). We defined three different errors between the predicted distributions and the ground truth exponential distributions (GT).

1. 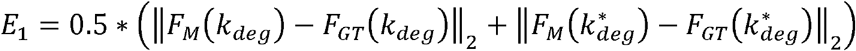 is the average L_2_ norm of the difference between the predicted cumulative distribution*F_M_*(·) and the ground truth *F_GT_*(·). This is the square root of the Cramér-von Mises metric.
2. 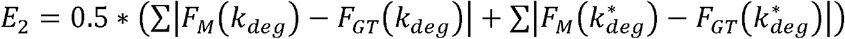 is the average sum of the absolute difference between the predicted cumulative distribution *F_M_*(·) and the ground truth *F_GT_*(·)
3. 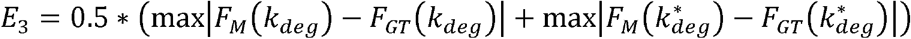 is the average of the maximum difference between the predicted cumulative distribution *F_M_*(·) and the ground truth *F_GT_*(·). This is the Kolmogorov-Smirnov metric.

The performance of DB and MERIDIAN based on these three errors can be found in Supplemental Information Table 4.

### Applying MERIDIAN to EGFR/Akt pathway

#### Inference of Lagrange multipliers from data is convex

The entropy functional

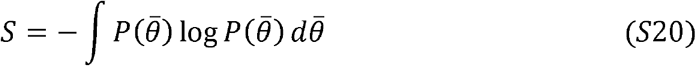

is convex with respect to the probability distribution 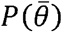. Moreover, the constraints that impose normalization and bin fractions are linear with respect to the probability distribution and are thus convex with respect to 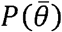 as well. Consequently, entire Lagrangian function (equation (4) of the main text)

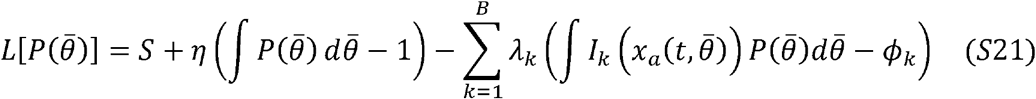

is also convex. Let us consider the dual problem in the space of Lagrange multipliers. We substitute the maximum entropy probability distribution 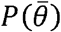 from equation (5) of the main text. We have the dual

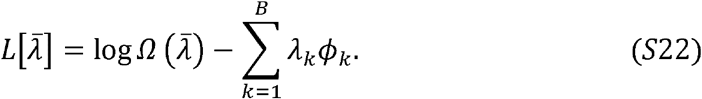

Given that the original objective function is convex, the maximization of the dual (equation S22) is equivalent to the problem of maximizing the original objective function (the entropy). Moreover, since the original problem is convex, the dual is convex as well (Bertsimas 1997). This can be easily checked because the Hessian of the Lagrangian function in equation (S22) is simply

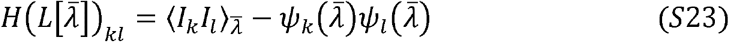

the covariance matrix between bin fraction indicator functions evaluated at 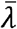. The covariance matrix is by construction a positive semi-definite matrix. This provides another evidence that the problem of finding Lagrange multipliers is a convex one.

#### Generalization of equation (5) for multiple species

Here, we give a generalization of equation (5) in the main text when the single cell distributions measured from multiple chemical species are used to constrain the parameter distribution. Consider that we have measured cell-to-cell variability in *n* different experimental conditions. The experimental conditions are identified by several indicators including identity of the measured species, input level, time of measurement, etc. We avoid multiple subscripts to specify these various indicators and denote the experimental conditions as {*x*^1^, *x*^2^,…, *x^n^*}. We consider that the single cell distribution at each measurement *“a”* is binned in *B_a_* bins. The maximum entropy parameter distribution is given by (see equation (5) in the main text)

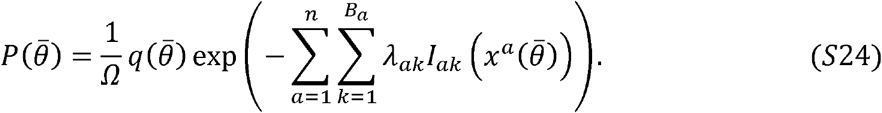

In equation (S24), *I_ak_*(*x*) is the indicator function corresponding to the *k*^th^ bin for the *a*^th^ experimental condition, *B_a_* is the number of bins representing the *a*^th^ experimental condition, and *λ_ak_* is the corresponding Lagrange multipliers.

#### Model of the EGFR/Akt signaling pathway

In this section, we describe in detail the dynamical model used to simulate levels of phosphorylated Akt as well as cell surface EGFRs after stimulation of cells with EGF.

The model of EGF/EGFR dependent phosphorylation of Akt was based on the previous work of Chen et al. (Chen, Schoeberl et al. 2009). We retained the branch of the Chen et al. model that leads to phosphorylation of Akt subsequent to EGF stimulation. The model had 17 species and 20 parameters. The description of the species is given in Supplemental information Table 3. The description of the parameters is given in Supplemental information Table 2. A system of ordinary differential equations describing dynamics of concentrations of species participating in signaling is given below (equations S24–S39). The model described EGF binding to EGFRs, subsequent receptors dimerization, phosphorylation, dephosphorylation, receptors internalization, degradation and delivery to cell surface and activation of Akt. We denote by active receptors phosphorylated receptors and by inactive receptors all other receptor states. In agreement with the literature only cell surface-localized phosphorylated receptors were allowed to activate Akt (Nicholson and Anderson 2002). We simplified the phosphorylation of pAkt through pEGFR; we implemented direct interaction between pEGFR and Akt leading to phosphorylation of Akt.

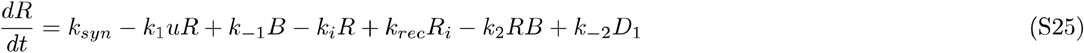

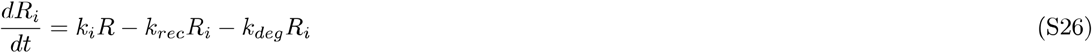

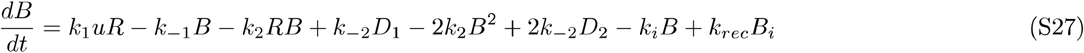

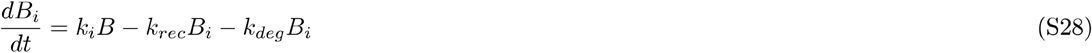

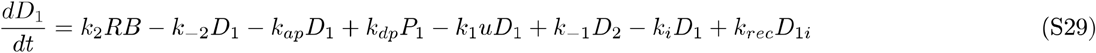

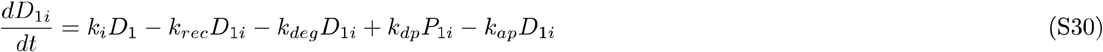

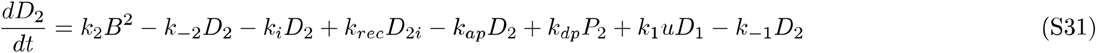

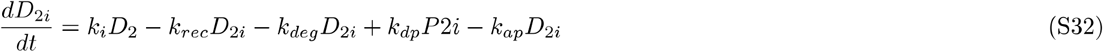

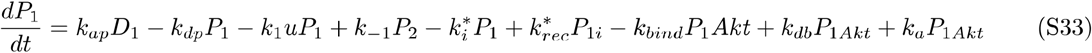

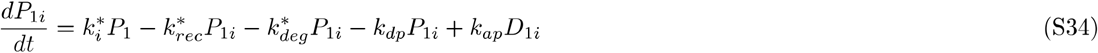

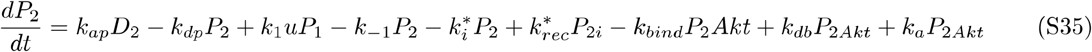

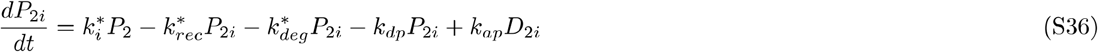

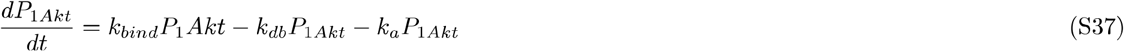

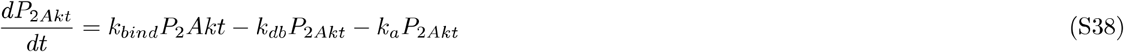

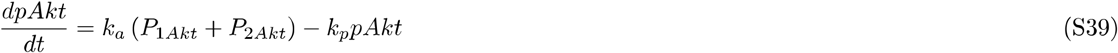

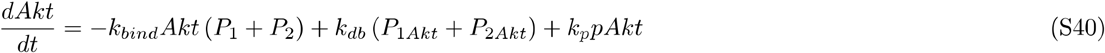

#### Binning single cell data

To infer the joint distribution over model parameters, we used 24 measured distributions of cell-to-cell variability (20 pAkt distributions, 1 pAkt background fluorescence distribution and 3 sEGFR distributions, see below). For each measured distribution we used 11 bins. The locations and widths of the bins were chosen to fully cover the observed abundance range while also ensuring reliable estimates of the bin fractions 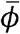. See Supplementary Information Table 1 for bin locations and experimentally estimated bin fractions.

We detected a small but significant pAkt signal in the absence of EGF stimulation. This background fluorescence signal likely originated from off target binding of pAkt-detecting antibodies. We assumed that the fluorescent readouts of pAkt/sEGFR levels in individual cells were equal to the sum of EGF dependent pAkt/sEGFR levels as computed using the signaling network model and the cell-dependent, but time-independent background fluorescence signal. In case of pAkt levels, the distribution of the background fluorescence was fitted to the experimentally measured distribution of the background fluorescence (pAkt readout without EGF stimulation). Unlike pAkt levels that respond to stimulation with EGF, cells maintain a high number of EGF receptors on the cell surface in the absence of EGF. As a result, we did not have experimental access to ‘background fluorescence’ distribution for sEGFR-detecting antibodies. We determined the range of background sEGFR fluorescence levels as follows. At the highest saturating dose of EGF (100 ng/ml) majority of the cell surface EGFRs are likely to be removed from the cell surface and degraded. At this dose, we assumed that the sEGFR background fluorescence can account for half of the measured fluorescence. We did not fit the distribution of background sEGFR levels to a specific distribution.

#### Numerical inference of Lagrange multipliers

The numerical search for Lagrange multipliers that are associated with bin fractions is a convex optimization problem (see above). We resorted to a straightforward and stable algorithm proposed in (Tkacik, Schneidman et al. 2006). The algorithm proceeded as follows. We started the calculations with an initial guess for the Lagrange multipliers at zero for each of the 11 bins of the 24 fitted distributions. In the *n*^th^ iteration, using the Lagrange multipliers 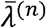, we estimated the predicted bin fractions 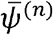 using Markov chain Monte Carlo (MCMC) sampling.

MCMC sampling in each iteration was performed as follows. We propagated 50 parallel chains starting at random points in the parameter space. Individual MCMC chains in the parameter space were run as follows. In the MCMC, on an average 10 parameters were changed in a single Monte Carlo step. The parameters were constrained to be within the upper and lower limits determined individually for each parameter based on available literature estimates (see Supplemental Information Table 2). Each chain was run for 50000 MCMC steps. At each step, we solved the system of differential equations given in equations (S25)–(S40) numerically with the proposed parameter assignment using the ode15s function of MATLAB. We evaluated the pAkt and sEGFR levels and accepted the proposed parameters using the Metropolis criterion applied to equation (5) in the main text. Briefly, for any set of parameters, we defined the energy

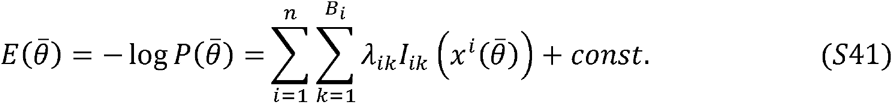

Starting from any parameter set 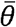, a new parameter set 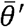 was proposed as described above. Then, the differential equations describing system dynamics were solved and the new energy 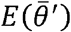 was computed. The difference in energy 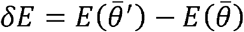 was used to probabilistically accept/reject the new parameter set with an acceptance probability

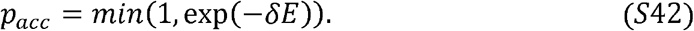

Parameter points that predicted pAkt and sEGFR levels outside of the ranges observed in experimental data were rejected (see Supplemental Information Table 5 for allowed ranges). We discarded the first 5000 steps as equilibration and saved parameter values every 50^th^ iteration. At the end of the calculation, parameter samples from all MCMC chains were combined together. We also imposed a few realistic constraints on pAkt and sEGFR time courses predicted by the model. All parameter sets that did not satisfy these constraints were discarded. The constraints were as follows. (1) Given that EGF ligand induces receptor endocytosis, we required that the surface EGFR levels at 180 minutes of sustained stimulation with 100 ng/ml EGF to be lower than the steady state surface EGFR levels in the absence of EGF stimulation. (2) Similarly we required that pAkt levels at 45 minutes were lower than pAkt levels at 5 minutes for the highest EGF stimulation (100 ng/ml).

Using the sampled parameters, we estimated the bin fractions 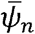 as well as the elements of the relative error vector 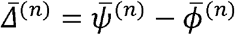 in the *n^th^* iteration.

For the *n*+1^st^ iteration, we proposed new multipliers 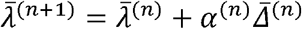. The multiplication constant *α*^(*n*)^ was chosen as follows. First, the approximate estimate of the predicted bin fractions for a given value of *α*^(*n*)^ was obtained using the Taylor series expansion

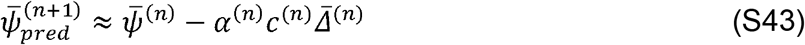

where *c*^(*n*)^ is the covariance matrix with entries

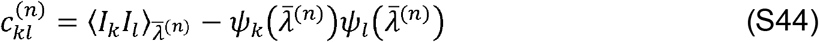

when the Lagrange multipliers are fixed at 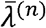. We chose *α*^(*n*)^ in the interval [0.05, A_*n*_] so as to minimize the predicted error 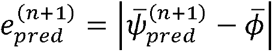.

1000 MC steps took 5-10 minutes. At the end of the calculation, the numerically inferred distribution over parameters captured with high accuracy the individual bin fractions of the distributions that were used to constrain it (Pearson’s *r*^2^ = 0.9, *p* < 10^−10^, median relative error = 14%). Notably, as seen in SI Figure 3, the predicted bin fractions from two independent calculations to determine the Lagrange multipliers were highly correlated with each other (Pearson’s *r*^2^ = 0.99, *p* < 10^−10^) indicating that the calculations converged to the same parameter distribution.

#### Inversion of covariance matrix (equation (S10))

In order to make predictions using MERIDIAN, we first sample several parameter points from the parameter distribution 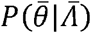 (equation (5) in the main text) using MCMC and the Metropolis criterion as described above. Using *N_S_* parameter samples, we generate a sparse matrix with entries *M_ab_* where *a* is the index of the sample point (*a* ∈ (0, *N_S_*)) and *b* is the index of the bin (and the experiment). There are a total of 24×11 = 264 bins used in this work and the *b* index runs between 1 and 264. The entry only if the model solutions pass through the *b*^th^ bin for any given set of parameters. From the matrix M, we estimate the 264×264 covariance matrix among the constraints. The entries of the convariance matrix are given by

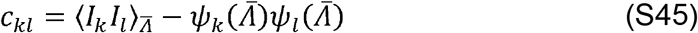

where *k,l* ∈ [1,264]. Next, we compute the inverse of the covariance matrix. Since all bin fractions at any given experimental conditions add up to one by definition, the covariance matrix is not full rank. Indeed, it has a total of 24 zero eigenvalues corresponding to 24 redundancies in the constrained single cell distributions. When inverting the covariance matrix, we neglect these 24 zero eigenvalues. The resultant inverse *c*^+^ is the so-called Moore-Penrose pseudoinverse of the matrix.

## SI Figures

**Supplemental Information Figure 1.**
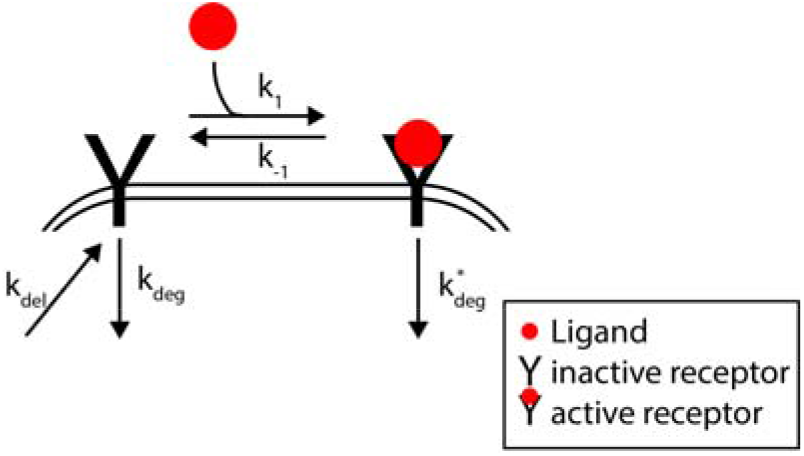
Schematic of the simplified growth factor model. Inactive receptors are delivered from the intracellular medium on the plasma membrane. Receptors bind to the ligand and get activated. Both activated and inactivated receptors are removed from the cell surface and degraded albeit at different rates.

**Supplemental Information Figure 2.**
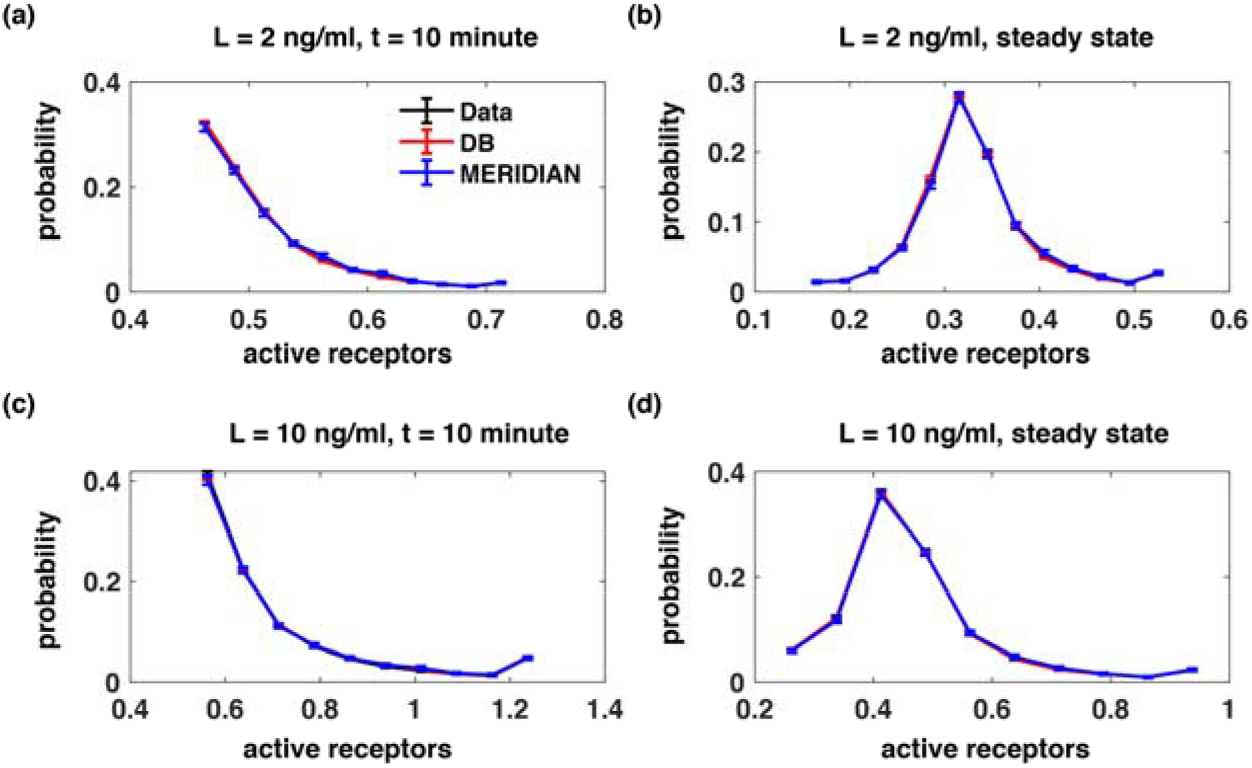
Fitted bin fractions using DB and MERIDIAN. We show the “ground truth” bin fractions (black lines) and the corresponding fits using MERIDIAN (blue lines) and the DB approach (red lines) across four simulation conditions.

**Supplemental Information Figure 3.**
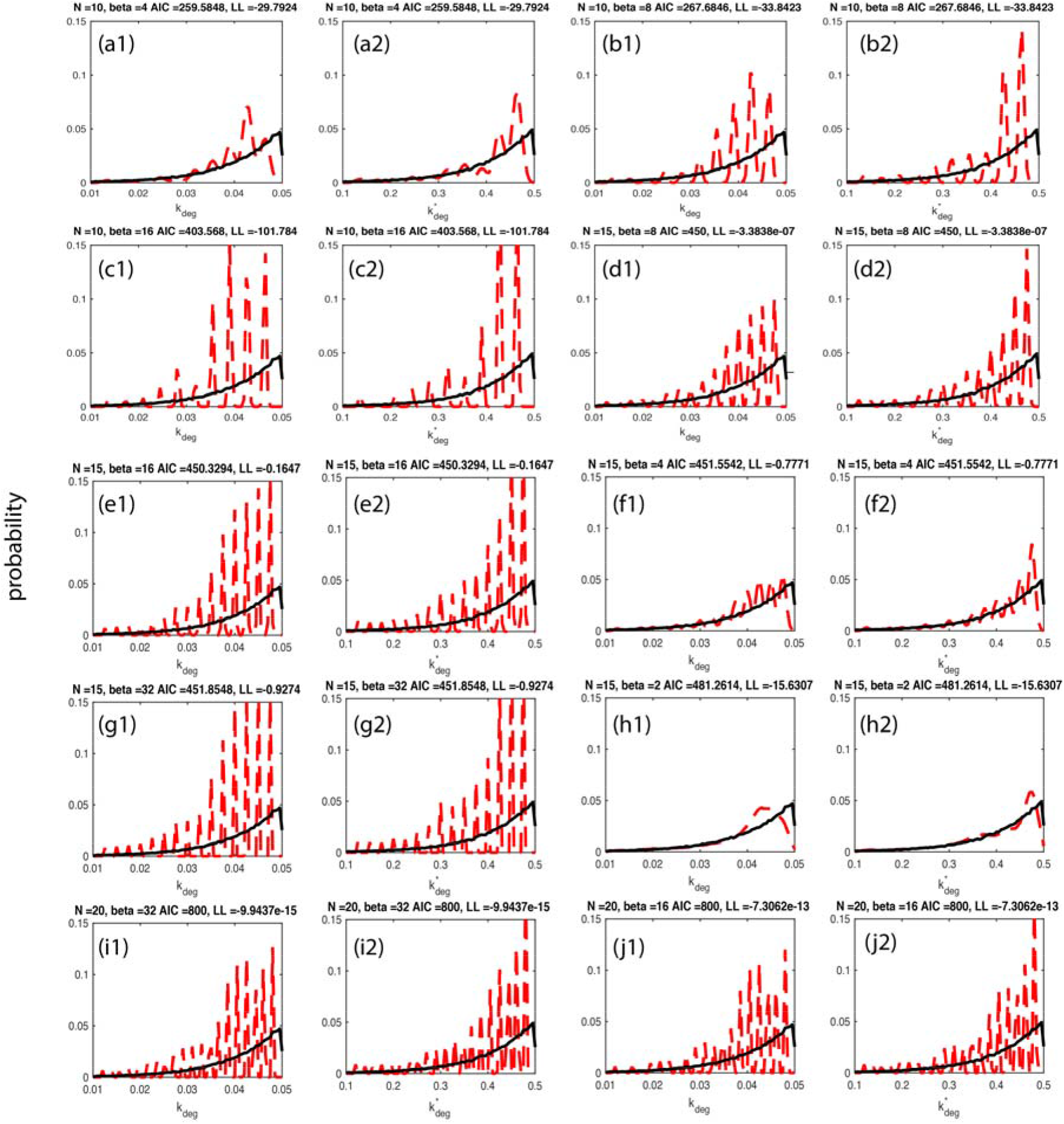
Inferred parameter distribution for top 10 DB fits. Panels (a1-j1) show the inferred distribution *P*(*k_deg_*) and panes (a2-j2) show the inferred distribution 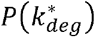 for top ten choices of method-parameters *N* (number of grid points) and *β* (width of the Gaussian basis function) according to the Akaike information score (dashed red line). The Akaike score and the likelihood are shown on the top of each panel. Black lines indicate the “ground truth” distributions.

**Supplemental Information Figure 4.**
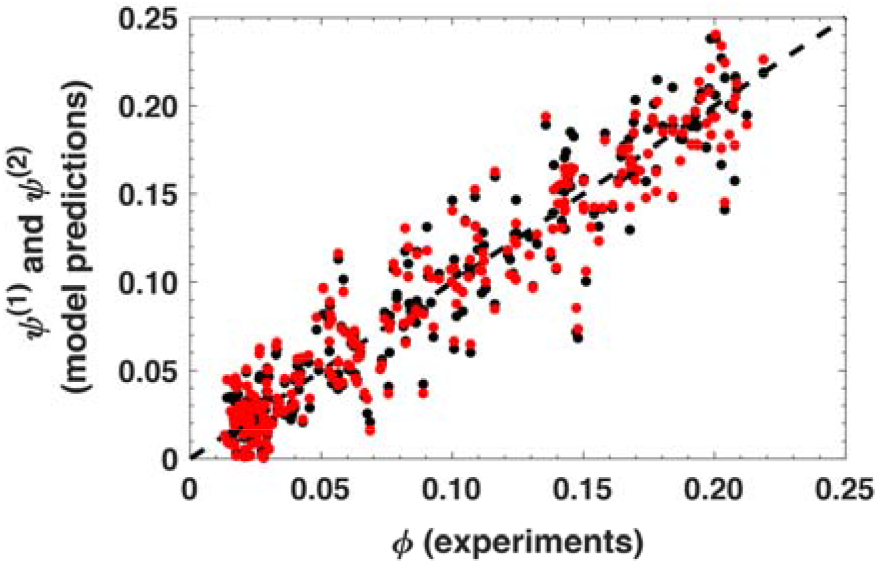
Model predictions agree for two independent calculations. The correlation between experimentally estimated bin fractions (x axis) and predicted bin fractions (y axis) for two independent searches for the Lagrange multipliers (red and blue dots).

**Supplemental Information Figure 5.**
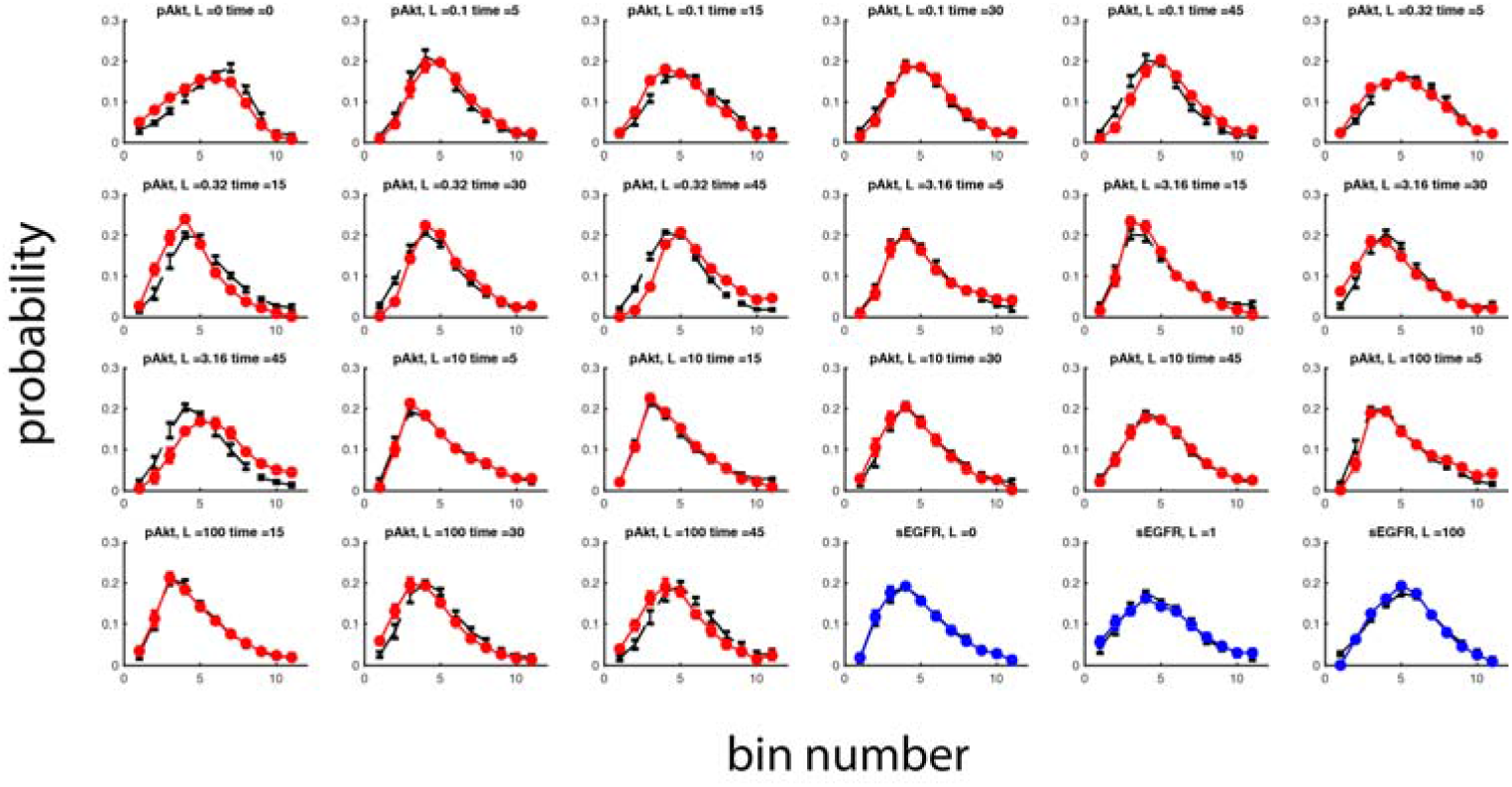
21 fitted pAkt distributions and 3 fitted sEGFR distributions used in parameter inference. We show all fitted single cell distributions (pAkt fits in red, sEGFR fits in blue, experimental data in black) used in the inference. Error bars represent experimentally estimated standard errors in the mean and model estimated uncertainties.

**Supplemental Information Figure 6.**
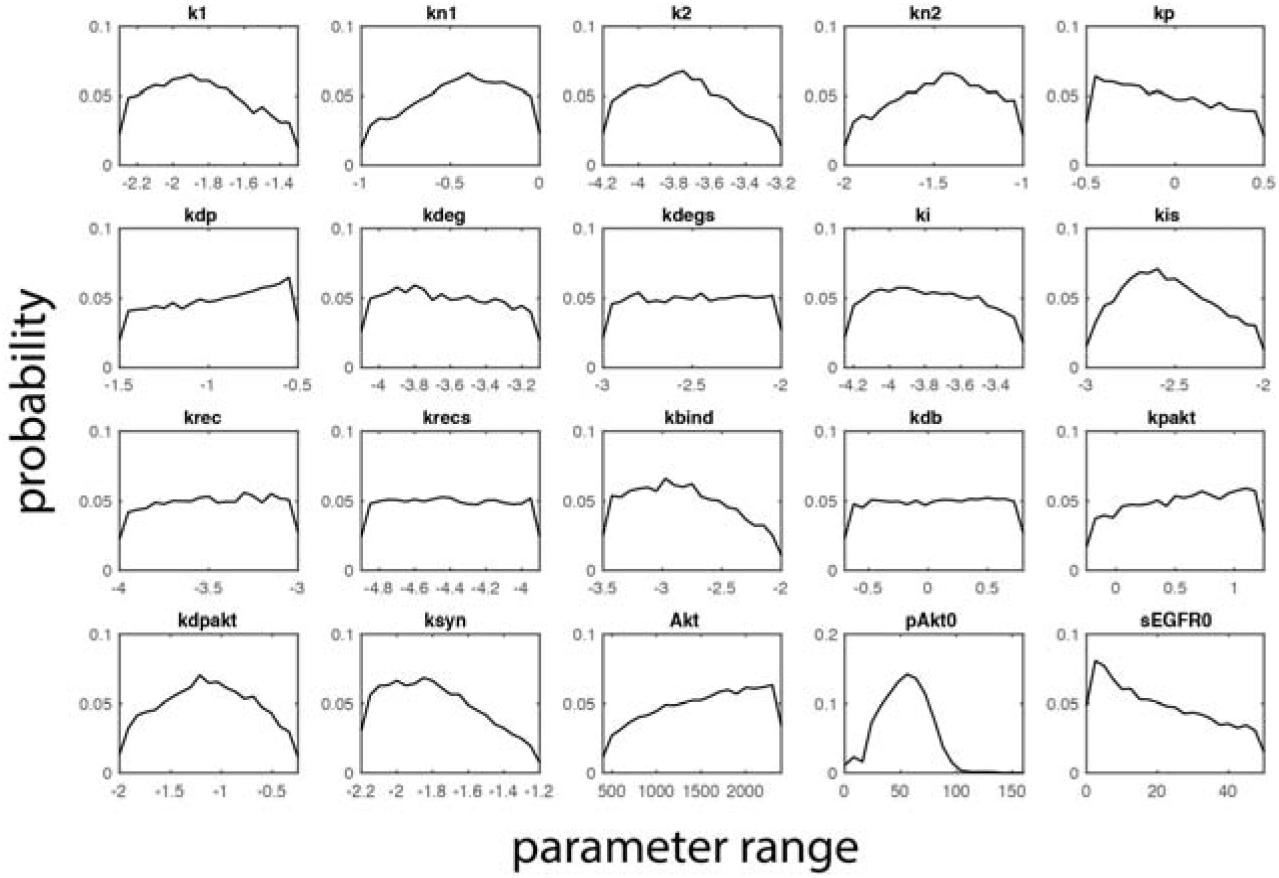
Inferred marginal distributions of all 20 model parameters.

**Supplemental Information Figure 7.**
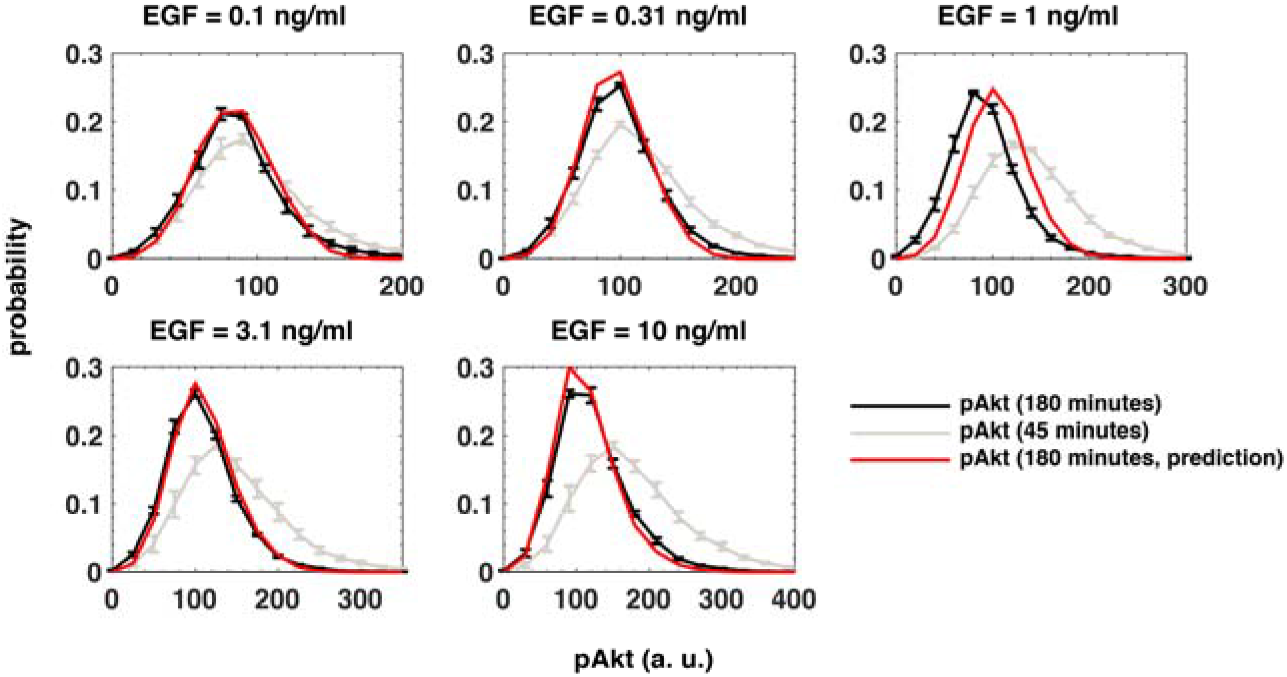
Population dynamics has not reached steady state at 45 minutes. We plot single cell distributions of pAkt levels at 45 minutes (gray lines) and at 180 minutes (experiments in black lines, model fit in red lines) across a broad range of EGF doses. Error bars represent experimentally estimated standard errors in the mean. For simplicity, we do not show the model estimated uncertainties.

